# Stem cell function *in vivo* is supported by an alternative glycolysis endpoint

**DOI:** 10.64898/2026.03.30.715412

**Authors:** Edward O. Kwarteng, Yafeng Li, Dieu Linh Nguyen, Michalis Agathocleous

## Abstract

Carbohydrates are classically catabolized by fermentation or oxidation, a choice that impacts many cellular functions including proliferation. Proliferating cells including somatic stem and progenitor cells are thought to favor fermentation over oxidation, and most proliferating cells *in vitro* depend on lactate production. However, it has not been tested if fermentation and oxidation are the universal obligatory terminal fates for carbohydrates *in vivo* because the key enzymes, lactate dehydrogenase (LDH) and pyruvate dehydrogenase (PDH), have not been simultaneously deleted in any cell type. Here we show that both fermentation and oxidation are dispensable for the survival and function of hematopoietic stem cells (HSC). Combined LDHA and LDHB deletion to ablate LDH did not impair HSC function, suggesting that HSCs and rapidly proliferating hematopoietic progenitors surprisingly do not require fermentation. Combined LDHA, LDHB, and PDH deletion abolished both glucose oxidation and fermentation, but did not impair HSC function. Glycolysis was preserved, suggesting the operation of an alternative endpoint. LDH/PDH-deficient HSCs terminated glycolysis through pyruvate export. Pyruvate export by HSCs and progenitors was a physiological response to changing nutrient levels. Quadruple deletion of LDHA/B, PDH, and the pyruvate transporter MCT1 impaired HSC function. This suggested that an essential role of glycolysis termination is not to produce acetyl-CoA or lactate but to remove pyruvate. Therefore, in contrast to classical theories and to *in vitro* metabolism, carbohydrate metabolism *in vivo* does not require oxidation or fermentation but can terminate directly in pyruvate export, and this alternative pathway is sufficient to support stem cell function.

## Introduction

Carbohydrate catabolism through glycolysis is a universal metabolic process that serves energy production, redox balance, and biosynthesis^1^. Classically glycolysis terminates by conversion of pyruvate either to lactate by LDH or to acetyl-CoA by PDH and then to CO_2_ through the tricarboxylic acid (TCA) cycle^1^ (**Figure 1A**). The idea that cells catabolize carbohydrates through either oxidation or fermentation was formulated through mainly *in vitro* work showing that cells switch between these two routes depending on environmental conditions such as oxygen availability^2^. This is the earliest known^3,4^ and most well studied metabolic choice. It is an underlying premise of much work on central carbon metabolism which assumes that inhibiting fermentation switches cells to oxidation and vice-versa. However, LDH and PDH have not been simultaneously deleted in any cell type to test if oxidation and fermentation are the obligatory terminal fates for carbohydrates *in vivo*, or if cells can sustain their metabolism and function without either LDH or PDH.

**Figure 1.**
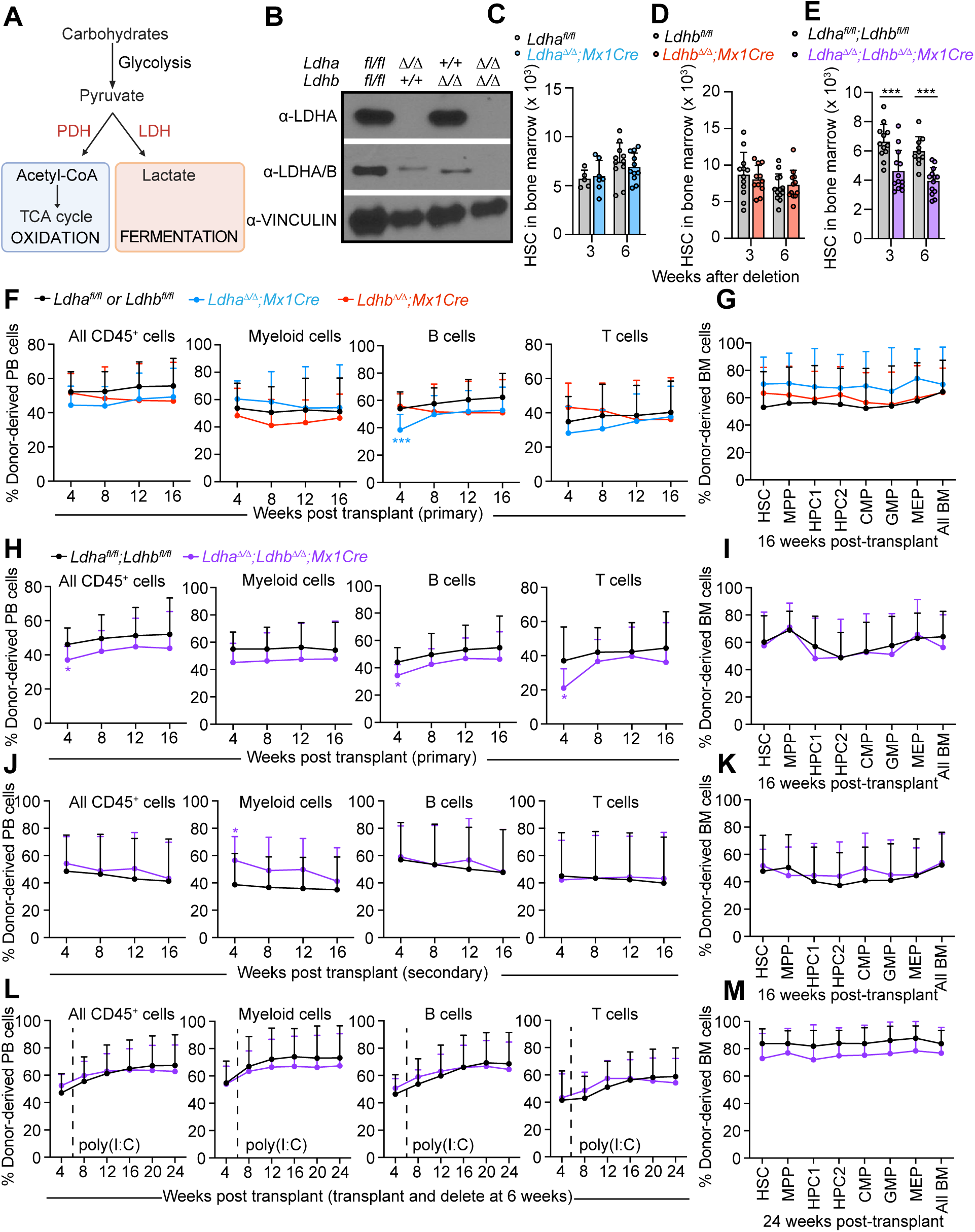
Glycolytic lactate production is dispensable for HSC survival and function. **A,** Schematic of canonical glycolysis terminating via fermentation of glucose to lactate by lactate dehydrogenase (LDH) or via conversion of pyruvate to acetyl-CoA by pyruvate dehydrogenase (PDH) for oxidation in the tricarboxylic acid (TCA) cycle. **B,** Immunoblot showing loss of LDHA and LDHB proteins in bone marrow cells following ACK lysis to remove red blood cells, 3 weeks after *Mx1Cre* induction with poly(I:C). **C–E,** The number of bone marrow HSCs in two legs in *Mx1Cre;Ldha^Δ/Δ^* (c), *Mx1Cre;Ldhb^Δ/Δ^* (**D**), and *Mx1Cre;Ldha^Δ/Δ^ ;Ldhb^Δ/Δ^* (**E**) mice and littermate controls at 3 and 6 weeks after deletion. **F–G,** Competitive BM transplantation of donor CD45.2^+^ *Mx1Cre;Ldha^Δ/Δ^*, *Mx1Cre;Ldhb^Δ/Δ^*, or *Ldha^fl/fl^ or Ldhb^fl/fl^* littermate control bone marrow cells with wild-type CD45.1;CD45.2 competitor cells to lethally irradiated CD45.1 recipient mice. Shown are donor chimerism in **F** peripheral blood and **G** BM HSPCs (n = 12–15 mice per genotype for blood; n = 10–14 for BM; three independent experiments). **H–I,** Competitive transplantation of *Mx1Cre;Ldha^Δ/Δ^;Ldhb^Δ/Δ^* or littermate control BM cells. Donor contribution to **H** peripheral blood over 16 weeks and **I** BM HSPCs at week 16 (n = 11–15 for blood; n = 10–14 for BM; three independent experiments). **J–K,** Secondary transplantation of 5 × 10⁶ BM cells from primary recipients shown in **H-I**. Donor chimerism in **J** peripheral blood and **K** BM HSPCs (n = 13–14 from three independent experiments). **L–M,** Competitive transplantation of undeleted donor *Mx1Cre;Ldha^fl/fl^;Ldhb^fl/fl^*or *Ldha^fl/fl^*;*Ldhb^fl/fl^* littermate control bone marrow cells followed by deletion at 6 weeks. Donor chimerism in **l** blood and **M** BM HSPCs (n = 11–15 from three independent experiments). Data represent mean ± s.d. Statistical significance was assessed with a 1-way ANOVA (**F-G**) followed by Dunnett’s test, or with a t-test (rest). All figures show * p<0.05, ** p<0.01, ***p<0.001.

The choice of carbohydrate oxidation or fermentation is not simply a housekeeping decision as it impacts cell proliferation, cell fate, and cancer cell growth. High LDH activity is thought to be a cardinal feature of rapidly proliferating cells^2^, including stem and progenitor cells^5–13^ and cancer cells. LDH produces NAD^+^, which supports glycolysis and other reactions^2,12,13^, and lactate, which has signaling functions in proliferating cells^14^. PDH produces acetyl-CoA, which fuels the TCA cycle and other reactions. But cells can also use alternatives routes to produce acetyl-CoA and NAD^+^ and they can import lactate. This raises the question of why cells need to oxidize or ferment carbohydrates. Here, we test if PDH and LDH are essential *in vivo,* or if glycolysis can terminate via an alternative endpoint. We focused on hematopoiesis, the most proliferation-intensive process in the body^15^, as our prior work made it possible to analyze the metabolism of rare hematopoietic cell types including HSCs^16,17^.

### LDH is dispensable for HSC survival and function

In somatic cells L-lactate is produced by LDH enzymes composed of LDHA and/or LDHB subunits. LDHC is testis-specific^18^, and LDHD uses the minor enantiomer D-lactate^19^. Analysis of Depmap data^20^ showed that cells with low *Ldhb* expression were preferentially sensitive to *Ldha* deletion, and cells with low *Ldha* expression were preferentially sensitive to *Ldhb* deletion (**Figure S1A**). Therefore, most cell lines require either LDHA or LDHB, suggesting LDH is essential *in vitro*. Combined LDHA/LDHB deletion abolishes LDH activity and reduces proliferation in some cancer cells *in vitro*^21,22^ or T cells *in vivo*^23^ but LDHA/B have not been deleted together in stem or progenitor cells.

HSCs are thought to rely on glucose fermentation via LDH because they reside in the hypoxic bone marrow^9,10,24–34^. Consistent with this idea, PDH loss does not inhibit HSC function^16^, PDH activation by deletion of PDH kinases inhibits HSC function^10^, and LDHA deletion may reduce the ability of HSCs to reconstitute secondary transplant recipients^27^. Glycolysis increases when HSCs are activated^35^. HSCs express both LDHA and LDHB (**Figure S1B**). To test the role of LDHA in hematopoiesis, we conditionally deleted *Ldha* in young adult *Mx1Cre;Ldha^fl/fl^* mice using poly-inosine poly-cytidine (poly I:C) (**Figure S1C**). Efficient deletion was confirmed by loss of LDHA protein and LDH activity, and genotyping of HSC-derived colonies (**Figure 1B**, **Figure S1D-F**). *Ldha* deletion did not affect the number of bone marrow HSCs, multipotent progenitors (MPPs), or most restricted progenitors (**Figure 1C, Figure S1H-Q**). *Ldha^Δ/Δ^* mice had normal or increased bone marrow cellularity, white blood cell and platelet counts but became anemic, as expected from a requirement of LDHA in erythrocytes which lack mitochondria (**Figure S1R-U**), and consequently had elevated erythroid progenitors and reduced numbers of some myeloid and lymphoid cells (**Figure S1V-AB)**.

To test the role of LDHB in hematopoiesis, we generated *Mx1Cre;Ldhb^fl/fl^* mice. LDHB protein was efficiently depleted, and genotyping of colonies formed by *Ldhb^Δ/Δ^* HSCs confirmed 100% deletion (**Figure 1B**, **Figure S2A-C**). LDH activity in hematopoietic cells was unaffected by *Ldhb* deletion, consistent with LDHA being the predominant isoenzyme (**Figure S2D**). *Ldhb* deletion had no effect on the frequency or number of HSCs, MPPs, restricted progenitors, or mature cell types in the bone marrow or blood (**Figure 1D, Figure S2E-Y**). Therefore, *Ldhb* is dispensable for steady-state hematopoiesis.

We generated *Mx1Cre;Ldha^fl/fl^;Ldhb^fl/fl^* mice to completely eliminate LDH in hematopoietic cells (**Figure 1B**). Genotyping of *Ldha^Δ/Δ^*;*Ldhb^Δ/Δ^*HSC-derived colonies confirmed 100% deletion (**Figure S3A-B**). *Ldha^Δ/Δ^*;*Ldhb^Δ/Δ^* mice had a mildly hypocellular bone marrow with reduced HSCs, MPPs, and most restricted progenitors (**Figure 1E, Figure S3C-M**). *Ldha^Δ/Δ^;Ldhb^Δ/Δ^* mice had normal white blood cell and platelet counts but became anemic (**Figure S3N-P**). As expected from their anemia, these mice exhibited increased bone marrow erythroid progenitors and reduction of some myeloid and lymphoid cells (**Figure S3Q-W**).

To test if *Ldha* and/or *Ldhb* are cell-autonomously required for HSC function, we performed competitive transplants of bone marrow cells into irradiated recipient mice **(Figure S4A).** *Ldha* or *Ldhb* deletion did not affect long-term multilineage reconstitution of the blood or HSCs and progenitor reconstitution of the bone marrow in primary, secondary, or in the case of *Ldha,* tertiary transplant recipients (**Figure 1F-G, Figure S4)**. Therefore, neither *Ldha* nor *Ldhb* are cell-autonomously required for HSC function, myelopoiesis, or B lymphopoiesis. To assess if total LDH is required for stem cell function, we competitively transplanted *Ldha^Δ/Δ^;Ldhb^Δ/Δ^* bone marrow cells into irradiated recipients. Long-term multilineage reconstitution in the blood and HSC and progenitor reconstitution in the bone marrow of primary and secondary transplant recipients were unaffected by *Ldha;Ldhb* deletion (**Figure 1H-K**). To avoid non-cell-autonomous effects of LDH deletion on HSPCs before transplant, we competitively transplanted undeleted *Mx1Cre;Ldha^fl/fl^;Ldhb^fl/fl^*or littermate control bone marrow cells into irradiated mice, and deleted *Ldha;Ldhb* 6 weeks later (**Figure S4B**). *Ldha;Ldhb* deletion did not affect long-term multilineage reconstitution of the blood or HSPC reconstitution in the bone marrow (**Figure 1L-M**). These results suggest that, unexpectedly^2,5,36^, and in contrast to its requirement for proliferation in cell lines *in vitro* (**Figure S1A**), LDH is not required for rapid cell proliferation during hematopoietic regeneration *in vivo*. HSCs do not rely on fermentation for their survival, proliferation, self-renewal, differentiation to myeloid and lymphoid cells, and ability to regenerate the hematopoietic system.

### LDH deletion does not block glycolytic and oxidative metabolism in HSCs and progenitors

Metabolomics analysis showed that *Ldha;Ldhb* deletion changed the levels of a quarter of the detected metabolites in bone marrow cells (**Figure 2A-B, Figure S5A, Table S1**). Lactate levels declined and levels of pyruvate and its transamination product alanine increased (**Figure 2C**), indicating that net LDH flux in the marrow is in the pyruvate-to-lactate direction. Glycolytic intermediates were elevated, and TCA cycle-related metabolites were unchanged or modestly elevated by LDH deficiency (**Figure 2C**). Consistent with LDH regenerating NAD^+^, LDH deficiency decreased the NAD⁺/NADH ratio (**Figure 2C**). To determine how glucose is utilized in the absence of LDH, we traced U^13^C-glucose in mice. LDH deletion increased both absolute levels and glucose-derived labeling in glycolytic and some TCA cycle-related metabolites in hematopoietic cells (**Figure 2D-G, Figure S5B-C**). These results suggested that LDH deletion did not inhibit glycolysis and increased the contribution of glucose to glycolytic and TCA cycle metabolites.

**Figure 2.**
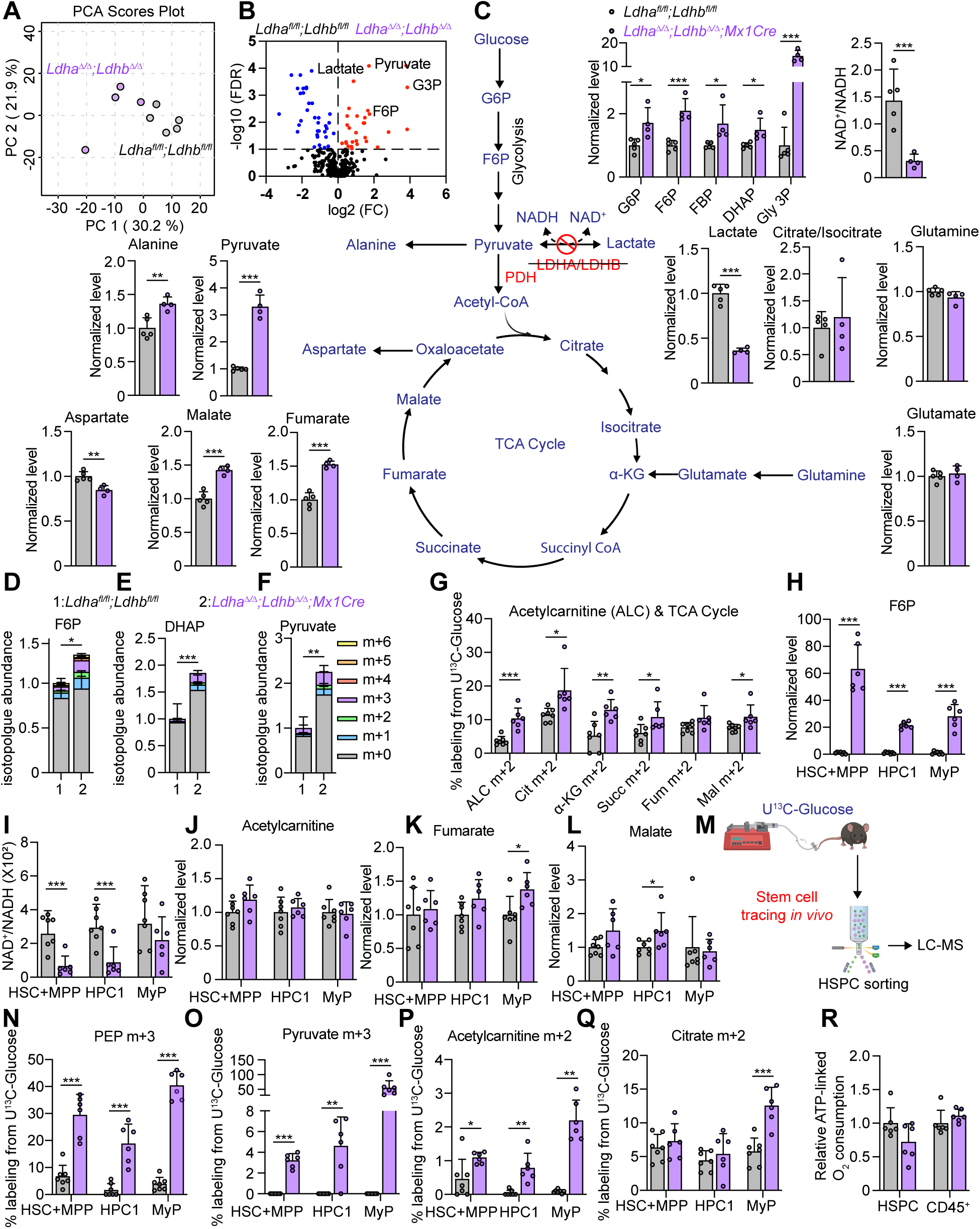
LDH deletion does not block glycolysis or oxidative 633 metabolism in HSCs progenitors. **A-C,** Metabolomics of bone marrow cells from *Mx1Cre;Ldha^Δ/Δ^*;*Ldhb^Δ/Δ^*mice or littermate controls 3 weeks after deletion. **A,** PCA scores plot of metabolomic profiles. **B,** Volcano plot of metabolite changes. Fold change represents metabolite levels in *Mx1Cre;Ldha^Δ/Δ^;Ldhb^Δ/Δ^* cells relative to controls. **C.** Levels of metabolites in central carbon metabolism. **D–F,** Isotopologue levels in indicated glycolytic metabolites after U^13^C-glucose tracing in *Mx1Cre;Ldha^Δ/Δ^*;*Ldhb^Δ/Δ^* mice and littermate controls (n = 6–7). **G,** Fractional enrichment of m+2 acetylcarnitine and TCA cycle intermediates after U^13^C-glucose tracing. **H–L,** Metabolite levels in the indicated HSC or progenitor populations. **M-Q,** *In vivo* U^13^C-glucose tracing in the indicated HSPC populations with fractional enrichment of phosphoenolpyruvate (PEP) m+3, pyruvate m+3, acetylcarnitine m+2 and citrate m+2. **R,** Respiratory oxygen consumption of sorted Lin^-^kit^+^ HSPCs and total CD45^+^ BM cells. All data represent mean ± s.d. Statistical significance was assessed with a Welch’s test (**H**) or Mann-Whitney test (**O-P**) or t-test (rest). Statistical significance for the NAD/NADH ratio was assessed by log transformation of data followed by unpaired t test. Statistical significance for metabolomics experiments was assessed with multiple t-tests controlling the false discovery rate at 5% with the Benjamini, Krieger, and Yekutieli method. All figures show * p<0.05, ** p<0.01, ***p<0.001.

To test the role of LDH in HSC metabolism, we performed rare cell metabolomics^17,37^. The levels of most detected metabolites are stable when HSCs and progenitors are isolated at cold temperatures^16,17,37^. We pooled CD150^+^CD48^-^Lin^-^Sca-1^+^Kit^+^ HSCs and CD150^-^CD48^-^Lin^-^Sca-1^+^Kit^+^ MPPs because these cell types are metabolically almost indistinguishable^17^. LDH deletion increased levels of detected glycolytic metabolites including fructose 6-phosphate, glucose 6-phosphate, and 3-phosphoglycerate, and decreased the NAD^+^/NADH ratio in HSC+MPP, CD150^-^CD48^+^Lin^-^Sca-1^+^Kit^+^ oligopotent hematopoietic progenitors (HPC1), and Lin⁻Sca1^-^Kit⁺ myeloid progenitors (MP) (**Figure 2H-I, Figure S5D-E, Tables S2-4**). LDH-deficient HSPCs showed no or minimal changes in TCA cycle-related metabolite levels (**Figure 2J-L**).

Studies of stem cell metabolism have typically relied on *in vitro* tracing^5^. To test how LDH deletion changes glucose use in HSCs and progenitors *in vivo*, we traced U^13^C-glucose in *Ldha^Δ/Δ^;Ldhb^Δ/Δ^* mice. LDH deletion increased glucose-derived labeling in glycerol-3-phosphate, PEP, pyruvate, and alanine in HSC/MPP, HPC1, or MP cells (**Figure 2N-O, Figure S5F-G**), suggesting LDH deletion increased or preserved glycolysis. Labeling of TCA cycle-related metabolites increased in some LDH-deficient HSPC types and did not change in others, consistent with increased or preserved glucose oxidation (**Figure 2P-Q, Figure S5H-J**). LDH deletion did not change respiratory O_2_ consumption in freshly sorted HSPCs and slightly increased it in CD45^+^ bone marrow cells **(Figure 2R)**. The combined results from metabolomics, stable isotope tracing, and O_2_ consumption indicate that LDH-deficient HSPCs surprisingly maintained or increased glycolytic flux, and that oxidative metabolism was also maintained or only slightly elevated.

### The impact of a complete oxidation and fermentation block on hematopoiesis

We previously showed that PDH deletion does not impair HSPC function^16^. To test if LDH-deficient HSPCs become reliant on PDH-mediated glucose oxidation, we tested the effects of combined inactivation of LDH and PDH on hematopoiesis. PDH and LDH have not been ablated together in any tissue to test if fermentation and oxidation are the sole and necessary endpoints of carbohydrate catabolism *in vivo*. *Ldha;Ldhb*-deficient mice were crossed to *Mx1Cre;Pdha1^fl^* mice (*Pdha1^Δ/Y^* males or *Pdha1^Δ/Δ^* females, here both denoted as *Pdha1^Δ^*) (**Figure S6A**). *Pdha1* encodes the essential PDH-E1α subunit of PDH, whose deletion ablates PDH activity^38^. *Mx1Cre* efficiently deletes *Pdha1* in HSCs^16^ and depletes PDH-E1α in hematopoietic cells (**Figure 3B**).

**Figure 3.**
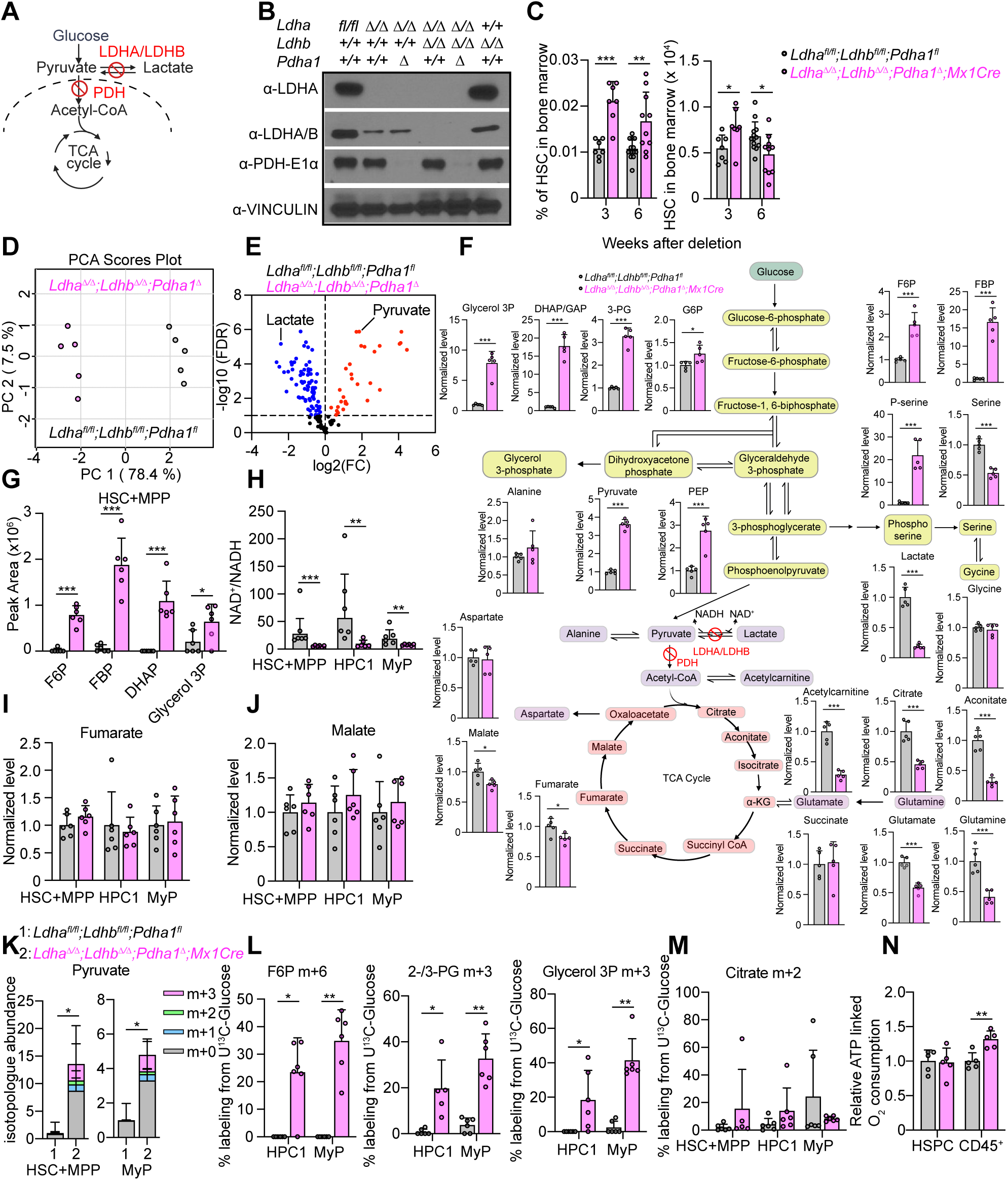
Carbohydrate oxidation and fermentation are dispensable for HSC and progenitor glycolysis, respiration, and survival. **A,** Schematic of combined LDH and PDH deletion. **B,** Immunoblot of LDHA, LDHB, and PDH in bone marrow cells following ACK lysis of red blood cells 3 weeks after deletion. **C,** HSC bone marrow frequency and number per two legs of mice from the indicated genotypes. **D-F,** Metabolomics of bone marrow cells from *Mx1Cre;Ldha^Δ/Δ^*;*Ldhb^Δ/Δ^*;*Pdha1^τι^* mice and littermate controls 3 weeks after deletion. **D,** PCA scores plot of the hematopoietic cell metabolome **E,** Volcano plot of metabolite changes in hematopoietic cells. Fold change represents metabolite levels in *Mx1Cre;Ldha^Δ/Δ^*;*Ldhb^Δ/Δ^*;*Pdha1^τι^* cells relative to controls. **F,** Levels of metabolites in central carbon metabolism. **G-J,** Levels of glycolytic, redox, and TCA cycle metabolites in HSC+MPP populations measured by *in vivo* metabolomics analysis. **K-M,** Results of *in vivo* U^13^C-glucose tracing in the indicated HSC and progenitor populations showing isotopologue abundance or fractional enrichment of the indicated metabolites. **N,** Respiratory ATP-linked O_2_ consumption of sorted Lin^-^kit^+^ HSPCs or total CD45^+^ bone marrow cells calculated as the difference in O_2_ consumption before and after oligomycin treatment, relative to control. Each point represents a biological replicate. All data represent mean ± s.d. Statistical significance was assessed with a Mann Whitney test (**G**, **K-L**) or a t-test (**C**, **F**, **N**). Statistical significance for the NAD^+^/NADH ratio was assessed by log transformation of data followed by a t test. Statistical significance for metabolomics experiments was assessed with multiple t-tests controlling the false discovery rate at 5% with the Benjamini, Krieger, and Yekutieli method. All figures show * p<0.05, ** p<0.01, ***p<0.001.

We first analyzed *Ldha^Δ/Δ^;Pdha1^Δ^* mice, which exhibited progressive reductions in white blood cells, red blood cells, and platelets (**Figure S6B-D**), defects more severe than in *Ldha^Δ/Δ^*or *Pdha1^Δ^* mice^16^. *Ldha^Δ/Δ^;Pdha1^Δ^* marrow cellularity and the number of neutrophils, monocytes, and B cells declined by 6 weeks post deletion (**Figure. S6E-I**). The number of Pre-CFU-E progenitors increased and the number of CFU-E progenitors and CD71^+^Ter119^+^ erythroblasts declined, in contrast to single knockouts, suggesting that CFU-Es require either LDHA or PDH (**Figure S6J-L**). Single *Ldha^Δ/Δ^;Pdha1^Δ^*HSCs sorted in methylcellulose formed much smaller colonies than *Ldha^Δ/Δ^*or *Pdha1^Δ^* HSCs^16^ (**Figure S6M-N**). This precluded HSC genotyping but suggested that 100% of HSCs were deleted. The number of *Ldha^Δ/Δ^;Pdha1^Δ^* HSCs and most multipotent and restricted progenitors did not change at 3 weeks and declined at 6 weeks after deletion (**Figure S6O-X**). The progressive reduction in hematopoietic cell numbers after LDHA;PDH deletion could be a consequence of either cell-autonomous or non-cell autonomous mechanisms, such as the severe anemia in these mice.

We generated *Mx1Cre;Ldha^Δ/Δ^;Ldhb^Δ/Δ^;Pdha1^Δ^*mice to completely block canonical glucose catabolic endpoints. Western blot confirmed loss of all three enzymes (**Figure 3A-B**). The frequency of *Ldha^Δ/Δ^;Ldhb^Δ/Δ^;Pdha1^Δ^* HSCs increased, and their number declined by 6 weeks after deletion (**Figure 3C**). The numbers of multipotent and restricted progenitors were mostly unchanged at 3 weeks and declined by 6 weeks, except for some megakaryocyte and erythroid lineage progenitors whose numbers increased (**Figure S7A-I**). Colony formation assays showed a severe reduction in the size of colonies from triple knockout HSCs, suggesting that 100% of HSCs were deleted, and that they require either LDH or PDH for their function *in vitro* (**Figure S7J**). Triple knockout mice exhibited bone marrow hypocellularity, severe anemia, leukopenia, and thrombocytopenia (**Figure S7K-N**). Erythroid development was arrested at the Pre-CFU-E to CFU-E stage, and this was accompanied by reduced numbers of neutrophils, monocytes and B cells (**Figure S7O-U**). This analysis suggested that, as expected from canonical depictions of glycolysis, erythroid progenitor development *in vivo*, and HSC colony formation *in vitro,* depend on the ability to switch between oxidation and fermentation. In contrast, the survival of HSCs and most restricted progenitors was not acutely affected by blocking oxidation and fermentation.

### HSCs and progenitors with combined LDH and PDH deletion maintain glycolysis and oxidative phosphorylation

To understand how cells metabolically function in the absence of canonical glycolysis termination, we performed metabolomics of triple knockout bone marrow cells. *Ldha^Δ/Δ^;Ldhb^Δ/Δ^;Pdha1^Δ^* cells were highly metabolically dissimilar from wild type cells with changes in the levels of most detected metabolites (**Figure 3D-F, Figure S8, Table S5**). Triple knockout cells had increased levels of glycolytic intermediates and pyruvate, and reduced lactate levels. The levels of many TCA cycle intermediates declined, consistent with a loss of carbohydrate oxidation (**Figure 3F**). To assess the fate of glucose in the absence of oxidation or fermentation we performed *in vivo* tracing with U^13^C-glucose. Triple knockout bone marrow cells had increased labeling of glycolysis intermediates, pyruvate (m+3), and alanine (m+3). Glucose-derived labeling in lactate (m+3), acetylcarnitine (m+2) and citrate (m+2) declined (**Figure S9A)**. These results suggested that concurrent deletion of LDH and PDH allows glucose to be metabolized to pyruvate which is neither converted to lactate nor oxidized in the TCA cycle.

To understand the metabolic effects of LDH and PDH deletion specifically in HSCs and progenitors, we used rare cell metabolomics. Triple knockout HSCs+MPPs, HPC1, and myeloid progenitor cells had increased levels of glycolytic intermediates, a reduction in NAD⁺/NADH and no changes in levels of detected TCA cycle metabolites (**Figure 3G-J, Figure S9B-C, Tables S6-8**). U^13^C glucose tracing *in vivo* showed that LDH/PDH deletion elevated both glucose-derived and non-glucose-derived pyruvate in HSCs and progenitors (**Figure 3K**). Glucose-derived labeling in glycolytic metabolites and alanine also increased, while labeling in citrate remained low in both wild type and LDH/PDH-deficient HSCs and progenitors (**Figure 3L-M, Figure. S9D**). Respiratory oxygen consumption was preserved in freshly sorted Lin^-^Kit^+^ HSPCs and slightly increased in CD45⁺ total bone marrow cells (**Figure 3M**), suggesting that in the absence of glucose oxidation, respiration is fueled by alternative substrates. These results suggested that glycolysis is maintained, and glucose-derived pyruvate accumulates in *Ldha^Δ/Δ^;Ldhb^Δ/Δ^;Pdha1^Δ^*HSPCs.

### Combined deletion of LDH and PDH does not impair HSC function

Regeneration of the ablated hematopoietic system imposes huge proliferative demands. To test if this process requires carbohydrate fermentation or oxidation, we transplanted *Ldha^Δ/Δ^;Pdha1^Δ^*, *Ldha^Δ/Δ^;Ldhb^Δ/Δ^;Pdha1^Δ^* and littermate control bone marrow with wild-type competitors into irradiated recipients (**Figure 4A-B**). *Ldha^Δ/Δ^;Pdha1^Δ^* and *Ldha^Δ/Δ^;Ldhb^Δ/Δ^;Pdha1^Δ^*donors showed normal myeloid and B cell reconstitution in the blood, and normal reconstitution of the HSC and progenitor compartments in the bone marrow (**Figure 4C-D)**. T cell reconstitution was impaired, consistent with our previous results showing that T cell development requires PDH^16^ **(Figure 4C).** Myeloid and B cell reconstitution and HSPC chimerism remained unaffected after serial transplantation, and T cell reconstitution deficits persisted (**Figure 4E-F**). To confirm that reconstituting donor HSCs had not escaped deletion, donor HSC-derived peripheral blood cells sorted from primary recipients were genotyped, showing efficient deletion of all three genes (**Figure S10**). These results suggested that the reduction in the numbers of HSCs, some progenitors, myeloid and B cells after *Ldha;Ldhb* or *Ldha;Ldhb;Pdha1* deletions were caused by non-cell autonomous mechanisms. Most importantly, these results suggested that HSC function *in vivo* – including survival, proliferation, self-renewal, multilineage differentiation, long-term production of mature cells, and the ability to regenerate the hematopoietic system after challenge – does not require LDH or PDH.

**Figure 4.**
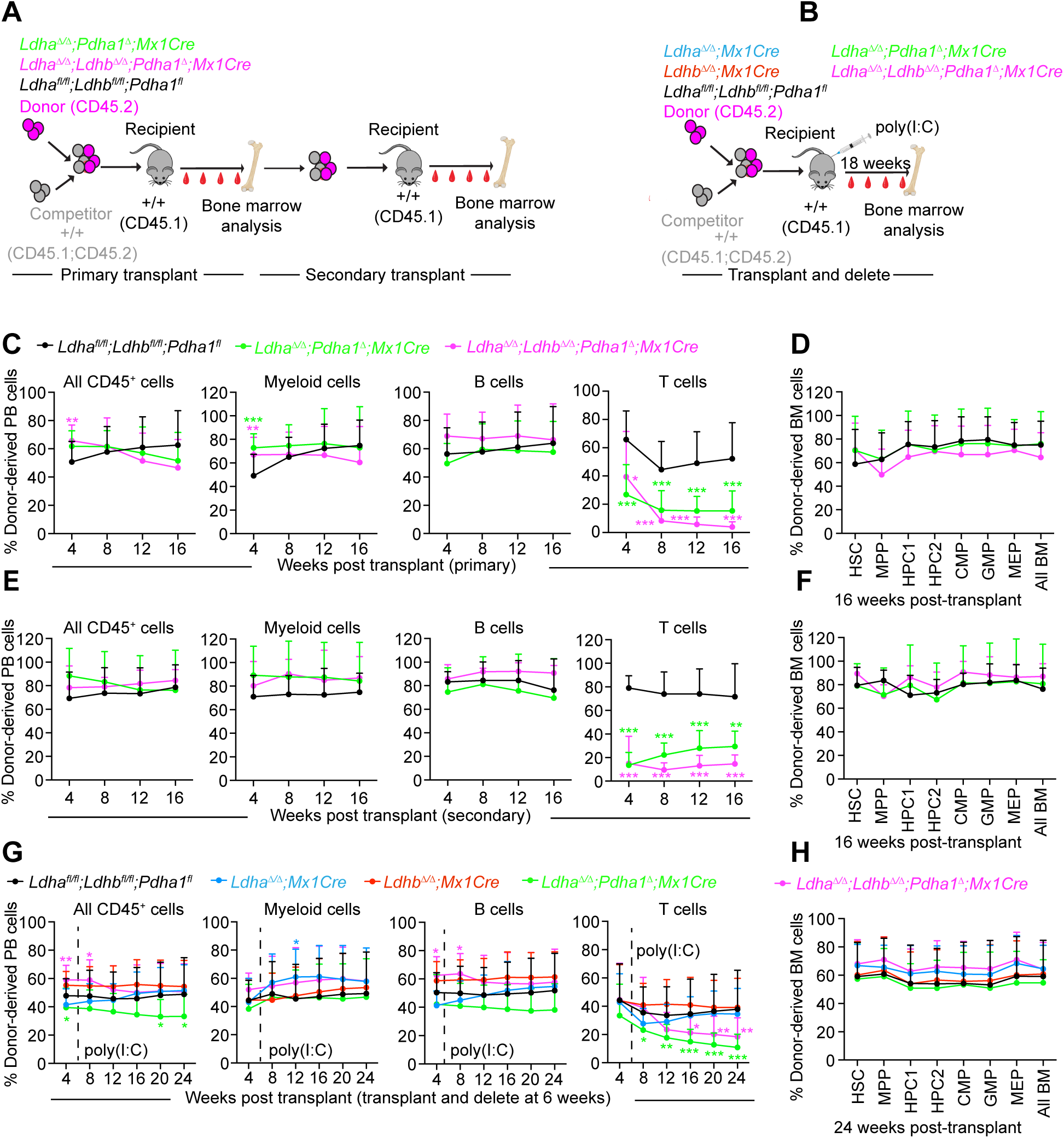
Combined ablation of carbohydrate oxidation and fermentation does not impair HSC function *in vivo*. Transplantation of *Mx1Cre;Ldha^Δ/Δ^*;*Ldhb^Δ/Δ^;Pdha1^τι^*, *Mx1Cre;Ldha^Δ/Δ^;Pdha1^τι^*, or littermate control CD45.2^+^ donor bone marrow cells with wild type CD45.1^+^CD45.2^+^ competitor cells into lethally irradiated CD45.1^+^ recipients. **A,** Schematic of primary and secondary transplantation experiments. **B,** Schematic of transplantation of undeleted donor cells followed by deletion at 6 weeks post-transplant. **C–D,** Donor chimerism in **C** blood and **D** bone marrow HSPCs in primary recipients (n = 12-13 for blood; n = 9-12 for BM from 3 independent experiments) from the transplant outlined in **A**. **E–F,** Donor contribution in **E** blood and **F** bone marrow HSPCs in secondary recipients (n = 8-11 for blood; n = 4-8 for BM from 3 independent experiments) from the transplant outlined in **A**. **G–H,** Peripheral blood (**G**) and BM HSPC (**H**) chimerism (n = 20–29 for blood; n = 13–24 for BM from 6 independent experiments) from the transplant outlined in **B.** All data represent mean ± s.d. Statistical significance was assessed with a 1-way ANOVA followed by Dunnett’s test. All figures show * p<0.05, ** p<0.01, ***p<0.001.

To rule out any non-cell autonomous effects of LDH and PDH deletion on HSCs before transplant, we competitively transplanted undeleted donor bone marrow cells from single, double, or triple knockout mice with wild type competitors, then deleted the floxed alleles in donor cells by treating recipient mice with poly I:C 6 weeks after transplant (**Figure 4B**). Myeloid and B cell chimerism were unaffected across genotypes (**Figure 4G**). T cell chimerism declined in *Ldha^Δ/Δ^;Pdha1^Δ^* and *Ldha^Δ/Δ^;Ldhb^Δ/Δ^;Pdha1^Δ^* donors (**Figure 4G**). *Ldha^Δ/Δ^;Pdha1^Δ^* and *Ldha^Δ/Δ^;Ldhb^Δ/Δ^;Pdha1^Δ^* cells did not significantly differ from control cells in their ability to reconstitute HSC and progenitor cell pools in the bone marrow (**Figure 4H**) Therefore, both carbohydrate oxidation and fermentation are dispensable for HSC function and for the production of myeloid and B cells.

### Glycolysis without oxidation and fermentation terminates principally in pyruvate export

We assessed the terminal fate of glucose in the absence of fermentation. Freshly sorted *Ldha^Δ/Δ^;Ldhb^Δ/Δ^* Lin⁻Kit⁺ HSPCs or total CD45^+^ bone marrow cells were incubated in culture for 12 hours, followed by metabolomics analysis of the medium to assess net nutrient consumption and metabolite production (**Figure 5A**). Wild-type HSPCs consumed less glucose and produced less lactate than wild-type CD45⁺ cells (**Figure 5B-C**). This is consistent with the functional data that HSPCs do not require LDH and suggests that HSPCs are not glycolytic. LDH deletion did not change the rate of glucose consumption but abolished lactate production and elevated pyruvate production. Metabolomic analysis indicated that pyruvate was the only detected metabolite overproduced by *Ldha^Δ/Δ^;Ldhb^Δ/Δ^*cells, and lactate was the main underproduced metabolite (**Figure 5D-E**). Lactate produced by wild type cells, and pyruvate produced by wild type or LDH-deficient cells originated predominantly from glucose (**Figure 5F-G**). These data suggest that HSPC glycolysis is maintained in the absence of LDH and terminates in pyruvate export.

**Figure 5.**
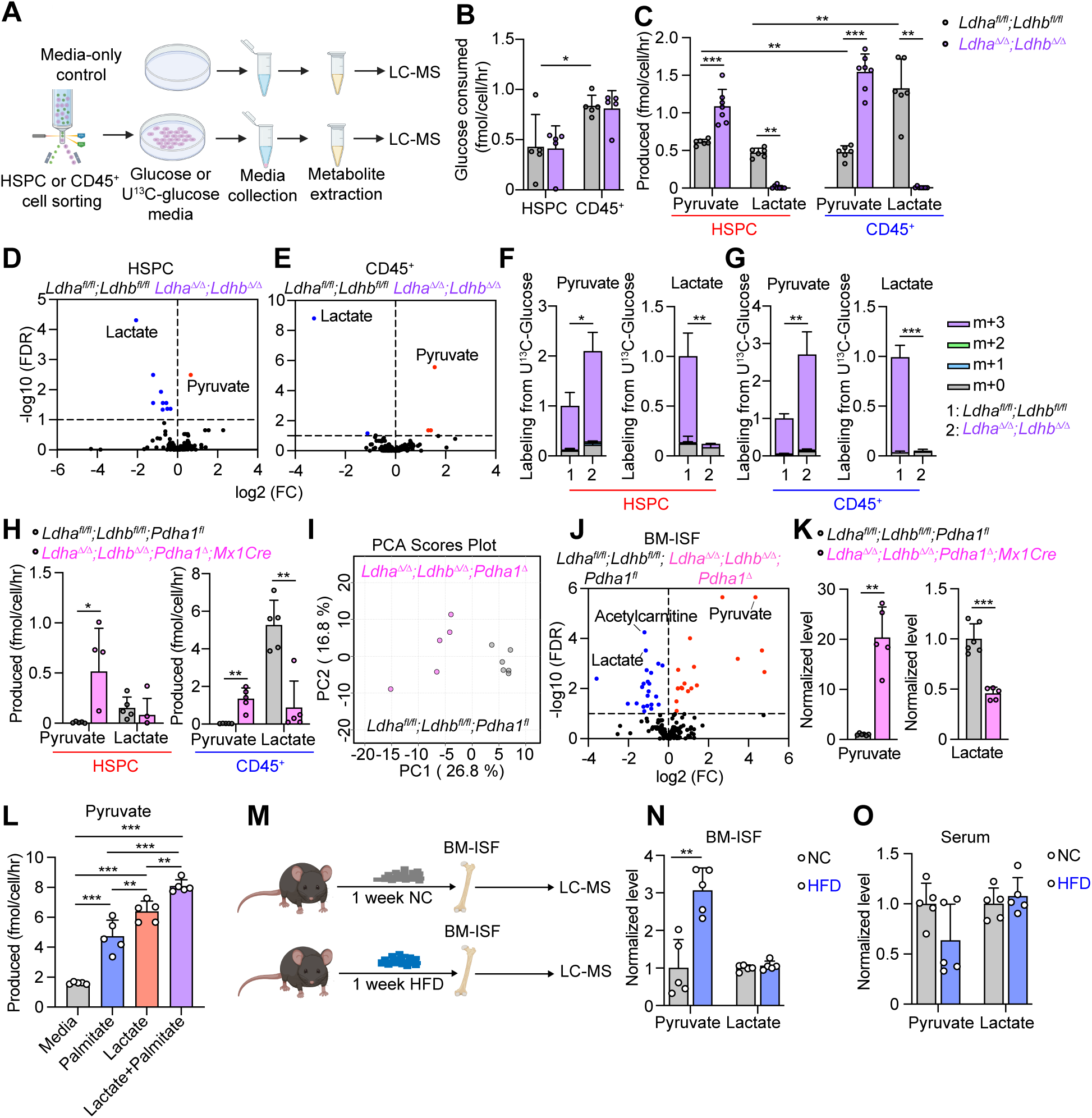
Glycolysis termination in pyruvate export. **A,** Schematic of glucose-consumption and metabolite-production assays. **B-C,** Glucose consumption and pyruvate and lactate production in sorted Lin^-^Kit^+^ HSPCs or CD45^+^ bone marrow cells after LDH deletion. Cells were cultured *ex vivo* with 0.5 mM glucose for 12 hours. depleted metabolite in sorted *Mx1Cre;Ldha^Δ/Δ^;Ldhb^Δ/Δ^* Lin^-^Kit^+^ HSPCs and CD45^+^ bone marrow cells. The x-axis shows fold change (FC) in *Ldha^Δ/Δ^;Ldhb^Δ/Δ^* relative to *Ldha^fl/fl^;Ldhb^fl/fl^* samples. **D-E,** Volcano plots showing pyruvate as the major elevated metabolite, and lactate as the major depleted metabolite in sorted *Mx1Cre;Ldha*D*/*D*;Ldhb*D*/*D Lin-Kit+ HSPCs and CD45+ bone marrow cells. The x-axis shows fold change (FC) in *Ldha*D*/*D*;Ldhb*D*/*D relative to *Ldhafl/fl;Ldhbfl/fl* samples. **F–G,** Glucose-derived pyruvate and lactate production in sorted HSPCs or CD45^+^ BM cells cultured for 12 hours with 0.5 mM U^13^C-glucose (n = 3–4 experiments). **H,** Pyruvate and lactate production in sorted Lin^-^Kit^+^ HSPCs or CD45^+^ bone marrow cells after LDH/PDH deletion. Cells were cultured *ex vivo* with 0.2 mM glucose for 12 hours. **I–J,** PCA plot (**I**) and volcano plot (**J**) of metabolites in BM interstitial fluid (BM-ISF). The x-axis in J shows fold change (FC) in *Ldha^Δ/Δ^;Ldhb^Δ/Δ^ ;Pdha1^Δ^* relative to *Ldha^fl/fl^;Ldhb^fl/fl^ ;Pdha1^fl^* samples. **K,** Pyruvate is highly enriched in the BM-ISF after triple deletion and lactate is highly depleted. **L,** Pyruvate production in sorted Lin^-^Kit^+^ HSPCs bone marrow cells cultured *ex vivo* with 1 mM glucose with or without 200 μM palmitate and 5 mM lactate for 12 hours. **M,** Schematic of BM-ISF metabolomics experiments in mice fed with high fat diet or normal chow for 1 week. **N–O,** Levels of pyruvate and lactate in **N**, BM-ISF and **O** serum after high fat diet or normal chow feeding. All data represent mean ± s.d. Statistical significance was assessed with a Mann-Whitney test (**C**-lactate), Welch’s test (**H**, **K**), or a t-test (rest). All figures show * p<0.05, ** p<0.01, ***p<0.001.

Changes in metabolic terminal fates *in vivo* should be reflected in the bone marrow interstitial fluid (ISF). Metabolites overproduced by LDH-deficient hematopoietic cells should be enriched and those underproduced should be depleted in the ISF. The top enriched metabolite in *Ldha^Δ/Δ^;Ldhb^Δ/Δ^*ISF was pyruvate, which accumulated 18-fold, and the top depleted metabolite was lactate (**Figure S11A-D, Table S9**). Tracing with U^13^C-glucose showed that glucose-derived pyruvate increased in *Ldha^Δ/Δ^;Ldhb^Δ/Δ^* ISF suggesting the elevated pyruvate partly originates from hematopoietic cell glycolysis (**Figure S11E-F**). The proportion of glucose-derived lactate decreased in *Ldha^Δ/Δ^;Ldhb^Δ/Δ^* ISF, suggesting that BM-ISF lactate normally partly originates from hematopoietic cell glycolysis via LDH (**Figure S11G**).

To assess the terminal fates of glucose in the absence of both fermentation and oxidation we measured metabolite production from freshly sorted *Ldha^Δ/Δ^;Ldhb^Δ/Δ^;Pdha1^Δ^* Lin⁻Kit⁺ HSPCs or CD45^+^ total bone marrow cells incubated for 12 hours in culture. These cells exported large amounts of pyruvate, with little to no lactate production (**Figure 5H**). Analysis of the *Ldha^Δ/Δ^;Ldhb^Δ/Δ^;Pdha1^Δ^* BM-ISF similarly showed a massive accumulation of pyruvate, accompanied by depletion of lactate (**Figure 5I-K, Table S10**). Stable isotope tracing showed that hematopoietic LDH/PDH deficiency increased U^13^C-glucose-derived ISF pyruvate and decreased U^13^C-glucose-derived ISF lactate, suggesting these ISF metabolites derived from hematopoietic cell glycolysis (**Figure S11H-I**).

These experiments showed that LDH/PDH-deficient HSPCs overproduce pyruvate. Cells typically favor lactate production in hypoxia, and pyruvate oxidation during increased energy needs in normoxia. We thus assessed if cells favor pyruvate production in a physiological setting. In contrast to most culture media, the circulation contains abundant lactate and fatty acids. We reasoned that extracellular lactate would suppress the forward LDH reaction and extracellular fatty acids would suppress the PDH reaction^39,40^, including in HSCs^41^, thus favoring pyruvate production. To test that, we measured pyruvate production from freshly sorted wild-type HSPCs in the presence of lactate and palmitate. Lactate and palmitate individually or in combination increased production of pyruvate (**Figure 5L**, **Figure S11J**). To test if increased pyruvate production also occurs in response to changes in nutrient availability *in vivo,* wild-type mice were fed a high-fat diet. This elevated pyruvate but not lactate levels specifically in the bone marrow ISF but not the serum (**Figure 5M-O, Figure S11K**), consistent with increased pyruvate production from hematopoietic cells.

In conclusion, metabolomics and U^13^C-glucose tracing in HSCs and other hematopoietic cells, in the ISF, and in culture media converge on the conclusion that in the absence of oxidation and fermentation, glycolysis terminates mainly in pyruvate export. This third way of terminating glycolysis also occurs normally in the physiological context of abundant extracellular fatty acids and lactate.

### Blocking pyruvate transport impairs *Ldha^Δ/Δ^;Ldhb^Δ/Δ^;Pdha1^Δ^* HSC function

Having established that LDH/PDH-deficient HSPCs export pyruvate as a terminal glycolysis step, we hypothesized that a major function of glycolysis termination is to remove pyruvate (**Figure 6A**). If this is true, then blocking pyruvate export should preferentially impair the function of LDH/PDH-deficient HSPCs. Transport of pyruvate and other monocarboxylates is mediated by MCT transporters. Of the major pyruvate transporters, MCT1 (*Slc16a1*) is the only one expressed at high levels in HSCs and other progenitors, while MCT2 is expressed at low levels and MCT3 and MCT4 are not expressed (**Figure S12A-B**). To test if LDH;PDH-deficient HSPCs are preferentially sensitive to MCT1 inhibition, we transplanted control, *Ldha^Δ/Δ^;Ldhb^Δ/Δ^*, or *Ldha^Δ/Δ^;Ldhb^Δ/Δ^;Pdha1^τι^* hematopoietic cells with wild type competitors into lethally irradiated recipients. When competitive hematopoiesis was established 6 weeks post-transplant, we treated mice with the MCT1 inhibitor AZD3965^42^ for 7 days (**Figure S12C**). MCT1 inhibition did not preferentially affect *Ldha^Δ/Δ^;Ldhb^Δ/Δ^*HSPC as compared to wild type HSPC function. However, MCT1 inhibition significantly depleted donor *Ldha^Δ/Δ^;Ldhb^Δ/Δ^;Pdha1^Δ^*myeloid and B cells in the peripheral blood and HSCs and progenitors in the bone marrow (**Figure 6B-C, Figure S12D**), suggesting *Ldha^Δ/Δ^;Ldhb^Δ/Δ^;Pdha1^τι^* HSCs and progenitors are more sensitive to MCT1 inhibition than wild-type HSCs and progenitors within the same environment.

**Figure 6.**
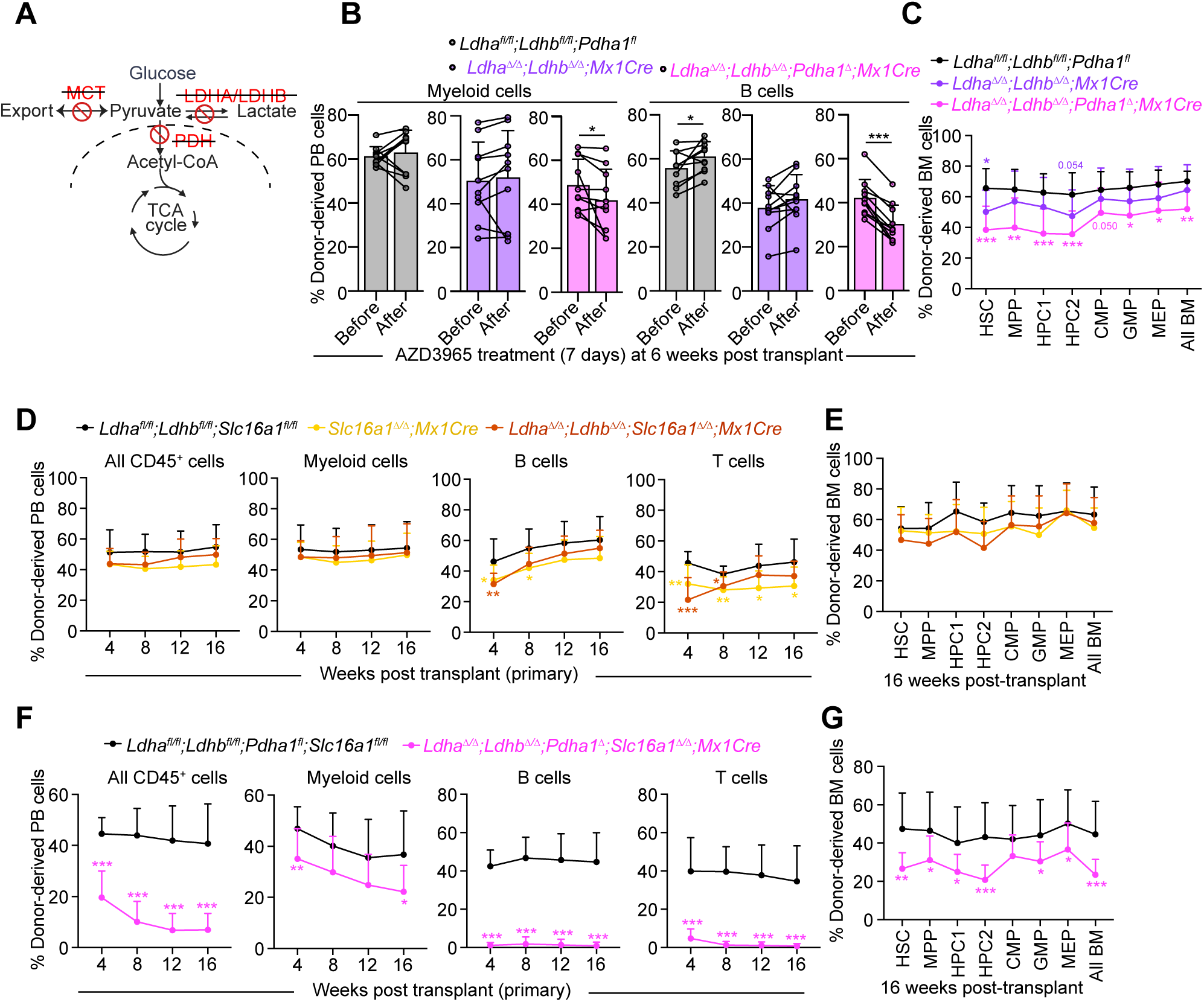
Deletion of MCT1 in the absence of LDH and PDH impairs HSC function. **A,** Schematic of glycolysis termination illustrating conditional *Ldha:Ldhb:Pdha1:Slc16a1* deletion with *Mx1Cre*. **B–C,** MCT1 inhibition with AZD3965 in chimeric transplant recipients reduces the fraction of donor *Ldha^Δ/Δ^;Ldhb^Δ/Δ^;Pdha1^Δ^* HSCs, progenitors, and mature cells: paired donor chimerism of **B** myeloid, B cells and **C** BM HSPCs (n = 10 mice from 3 independent experiments). **D–E,** Competitive bone marrow transplantation of donor CD45.2^+^ *Mx1Cre;Slc16a1^Δ/Δ^*, *Mx1Cre;Ldha^Δ/Δ^;Ldhb^Δ/Δ^*;*Slc16a1^Δ/Δ^*, or *Ldha^fl/fl^;Ldhb^fl/fl^*;*Slc16a1^fl/fl^* littermate control bone marrow cells with wild-type CD45.1;CD45.2 competitor cells to lethally irradiated CD45.1 recipient mice. Shown are donor chimerism in **D** peripheral blood at weeks 4-16 and **E** BM HSPCs at week 16 (n = 11–15 mice per genotype for blood; n = 12–15 for BM; three independent experiments). **F–G,** Competitive transplantation of *Mx1Cre;Ldha^Δ/Δ^;Ldhb^Δ/Δ^*;*Pdha1^Δ^*;*Slc16a1^Δ/Δ^* or littermate control BM cells. Donor contribution to **F** peripheral blood at weeks 4-16 and **G** BM HSPCs at week 16 (n = 13 for blood; n = 12–13 for BM; three independent experiments). Data represent mean ± s.d. Statistical significance was assessed with a paired t-test (**B**), one-way ANOVA (**C, D** - CD45^+^, Myeloid, B cells, **E**), a Brown-Forsythe ANOVA test (**D**: T cells), t-test (**F** – Myeloid cells), Mann-Whitney test (**F-** CD45^+^, B, T cells) and Welch’s test (**G**). All figures show * p<0.05, ** p<0.01, ***p<0.001.

Pharmacological MCT1 inhibition might incompletely block MCT1 or have unanticipated off-target effects, or effects in non-hematopoietic tissues. To test in a rigorous genetic way if pyruvate transport is essential specifically in LDH/PDH-deficient hematopoietic cells in a cell-autonomous manner, we generated quadruple *Mx1Cre;Ldha^fl/fl^;Ldhb^fl/f^;Pdha1^fl^;Slc16a1^fl/fl^*mice and their associated controls. Single *Slc16a1* deletion did not affect the number of HSCs or other bone marrow hematopoietic or blood cells 3 or 6 weeks after poly I:C and did not impair HSC function after competitive transplantation (**Figure S13**). Triple *Ldha;Ldhb;Slc16a1* deletion reduced numbers of HSCs, most progenitors and mature cells in the bone marrow and expanded erythropoiesis (**Figure S14**), effects similar to those of *Ldha;Ldhb* deletion (**Figure S3**). Competitive transplantation of *Slc16a1^Δ/Δ^*, *Ldha^Δ/Δ^;Ldhb^Δ/Δ^;Slc16a1^Δ/Δ^* or littermate control bone marrow with wild-type competitors into irradiated recipients revealed no defects in long-term multilineage reconstitution of leukocytes or hematopoietic stem and progenitor cells (**Figure 6D-E**). Quadruple *Ldha;Ldhb;Pdha1;Slc16a1* deletion caused widespread hematopoietic deficits including reduced numbers of HSCs and some progenitors, depleted neutrophils, B cells, and erythroid progenitors, leukopenia, anemia, and thrombocytopenia (**Figure S15)**. Competitive transplantation of *Ldha^Δ/Δ^;Ldhb^Δ/Δ^;Pdha1^Δ^;Slc16a1^Δ/Δ^*or littermate control bone marrow with wild-type competitors into irradiated recipients revealed profound deficits in multilineage reconstitution, with reduced donor-derived myeloid and lymphoid cells in the blood, and HSC and progenitor cells in the bone marrow (**Figure 6F-G**). Therefore, fermentation and oxidation are essential in HSCs only when pyruvate export is blocked.

## Discussion

Our work challenges the classical view that cells must either oxidize or ferment carbohydrates by demonstrating that LDH and PDH are dispensable for HSC function *in vivo*. Instead, HSCs can terminate glycolysis by exporting pyruvate, a mechanism sufficient to support their function at homeostasis and regeneration. Pyruvate production occurs not only after LDH;PDH deletion, but also in wild type cells in physiological conditions of high levels of extracellular lactate and palmitate. Although it has not been previously shown in any cell type that LDH and PDH are dispensable for cell survival and proliferation, breast cancer cells can export pyruvate through MCT1^43^ and muscle cells can export glycolysis-derived alanine^44^, suggesting glycolytic cells can use alternatives to fermentation and oxidation in pathophysiological settings.

The combined dispensability of LDH and PDH for HSCs and most multipotent and restricted myeloid progenitors is in contrast to their requirement in some erythroid and T cell progenitors and for HSC colony formation *in vitro*. Therefore, the classical view of glycolysis termination, determined largely from experiments *in vitro* or in bulk cell populations, holds in some specialized hematopoietic cells, but differs in HSCs and most hematopoietic progenitor cells *in vivo*.

Our results have implications for the link between glycolysis, cell proliferation, and somatic stem cells. Glycolysis terminating in high lactate production is a feature of many proliferating cells including stem or progenitor cells and cancer cells^2,5,36^. It is thought that this adaptation promotes cell proliferation^2,5,36^. Our results show that glycolytic lactate production is in fact not required for HSC function, or for myeloid progenitors, which are among the most proliferative cell types in the body^15^. High glycolytic production of lactate is therefore dispensable for both self-renewal and rapid cell proliferation. All three pyruvate disposal routes in HSCs must be blocked off through LDH, PDH, and MCT1 deletion in order to elicit phenotypic defects. This suggests that a key role of glycolysis termination *in vivo* is not production of acetyl-CoA via PDH or of lactate via LDH, but disposal of pyruvate via oxidation, fermentation, or export (**Figure S15L**).

## Methods

### Mice

*Ldha^fl/fl^* (Jackson Laboratory #030112)^27^, *Pdha1^fll^* (Jackson Laboratory #017443)^38^, *Slc16a1^fl^* (a gift from B. Morrison, Johns Hopkins)^45^ and *Mx1cre* (Jackson Laboratory #003556)^46^ mice were previously described. *Ldhb^fl/fl^* mice were generated by crossing *Ldhb^tm1a(KOMP)Wtsi/WtsiOulu^*mice (EMMA, EM:08936) with *Flpe* mice^47^. Mice were on a C57BL background. Mice were injected every other day with 40 μg poly (I:C) (Cytiva 274732-01) for a total of five intraperitoneal injections at 6-8 weeks to induce Cre expression. Mice were analyzed at 3 and 6 weeks from the first day of poly (I:C) injection. For experiments in which Cre expression were induced after transplantation, mice were injected every other day with 40 μg poly (I:C) for a total of seven injections. C57BL/Ka-Thy-1.1 (CD45.2) and C57BL/Ka-Thy-1.2 (CD45.1) mice were used for transplantation experiments. CD45.1;CD45.2 mice were used as competitors. Both male and female mice were used in all experiments. For high fat diet experiments, mice were fed normal chow (Teklad #2016, 12 kcal % from fat) or a high-fat diet (Research Diets D12492, 60 kcal % from fat) for 1 week. Mice were housed in the Animal Resource Center at the University of Texas Southwestern Medical Center. All procedures were approved by the UT Southwestern Institutional Animal Care and Use Committee.

### Hematopoietic analysis

For flow cytometry, bone marrow cells were obtained by flushing tibias and femurs of both legs with a 25G needle (analysis) or by crushing tibias, femurs, vertebrae and pelvic bones with mortar and pestle (for cell sorting or for chimerism analysis in transplants) in Ca^2+/^Mg^2+^ free Hank’s balanced salt solution (HBSS; Gibco) supplemented with 2% heat-inactivated bovine serum (HIBS; Gibco). Spleens were crushed and filtered through a 40 μm strainer (Thermo Fisher). Cells were counted with a Vi-CELL XR viability analyzer (Beckman coulter). Hematologic parameters were measured with an Element HT5 veterinary hematology analyzer (Heska corporation). Cells stained with fluorescent marker-conjugated surface antibodies for 30 minutes at 4°C for analysis of mature cell types, or for 90 minutes on ice for analysis of hematopoietic stem and progenitor cells (HSPCs). After incubation, cells were washed with HBSS + 2% HIBS and resuspend in HBSS + 2% HIBS containing propidium iodide or DAPI (1 μg/ml) for separation of live/dead cells. Cell populations were identified with the following markers^48–51^: CD150^+^CD48^-^Lineage^-^Sca-1^+^Kit^+^ hematopoietic stem cells (HSCs), CD150^-^CD48^-^Lineage^-^Sca-1^+^Kit^+^ multipotent progenitor cells (MPPs), CD150^-^CD48^+^Lineage^-^Sca-1^+^Kit^+^ (HPC1), CD150^+^CD48^+^Lineage^-^Sca-1^+^Kit^+^ (HPC2), CD34^+^CD16/32^-^Lineage^-^Sca-1^-^Kit^+^ common myeloid progenitors (CMPs), CD34^+^CD16/32^+^Lineage^-^Sca-1^-^Kit^+^ granulocyte-monocyte progenitors (GMPs), CD34^-^CD16/32^-^Lineage^-^Sca-1^-^Kit^+^ megakaryocyte-erythroid progenitors (MEPs), CD41^+^CD150^+^Lineage^-^Sca-1^-^Kit^+^ megakaryocyte progenitors (MkP), CD105^-^CD150^-^ CD41^-^CD16/32^-^Lineage^-^Sca-1^-^Kit^+^ pre-granulocyte-monocyte progenitors (Pre-GM), CD105^-^CD150^+^CD41^-^CD16/32^-^Lineage^-^Sca-1^-^Kit^+^ pre-megakaryocyte-erythroid progenitors (Pre-Meg-E), CD105^+^CD150^+^CD41^-^CD16/32^-^Lineage^-^Sca-1^-^Kit^+^ pre-colony forming unit-erythroid (Pre-CFU-E), CD105^+^CD150^-^CD41^-^CD16/32^-^Lineage^-^Sca-1^-^Kit^+^ colony forming unit-erythroid (CFU-E), Mac-1^+^CD115^-^Ly6G^+^ neutrophils, Mac-1^+^CD115^+^Ly6G^-^ monocytes, CD71^+^Ter-119^+^ erythroblasts, CD3^+^ T cells, and B220^+^ B cells. The Lineage antibody cocktail used for hematopoietic stem and progenitor cell (HSPC) identification consisted of CD2, CD3, CD5, CD8, Ter-119, B220, and Gr-1 antibodies. Flow cytometry antibodies were acquired from BD Biosciences, eBiosciences, BioLegend, or Tonbo biosciences. For isolation of HSPCs, cells were incubated with paramagnetic CD117 microbeads and enriched with a QuadroMACS magnetic separator (Miltenyi Biotec) before cell sorting. Analysis and cell sorting were performed using BD LSRFortessa (BD Biosciences), BD FACSCanto (BD Biosciences) or BD FACSSymphony S6 (BD Biosciences). Data were analyzed using Flowjo (Flowjo LLC).

### Bone marrow reconstitution assays

For competitive transplantation experiments, 500,000-1,000,000 cells each of donor (CD45.2) and competitor (CD45.1/CD45.2) cells were mixed and transplanted into the retro-orbital venous sinus of anesthetized CD45.1 recipients irradiated using an XRAD 320 X-ray irradiator (Precision X-Ray) with two doses of 540 rad (1080 rad in total) at least 3 hours apart. Donor cells were obtained from experimental or littermate control donor 6-8 weeks old mice 3 weeks after the start of poly I:C administration. In experiments in which deletion was induced after transplant, poly I:C was administered 6 weeks post-transplant. Recipient mice were maintained on antibiotic water (Baytril 0.08 mg/ml) for 1 week before and 4 weeks after transplantation. Peripheral blood was obtained from the tail veins of recipient mice every four weeks. Red blood cells were lysed with ammonium chloride potassium buffer. Samples were stained with CD45.2 (104), CD45.1 (A20), B220, Mac1, CD3, CD4, CD8, Ly6c, Ly6g, and CD115 antibodies, and analyzed with BD LSRFortessa (BD Biosciences). At the experimental endpoint, recipient mice were euthanized, and bone marrow cells was obtained from crushed femurs, tibias, pelvic bones, and spine for surface antibody staining and flow cytometry analysis or for serial transplants. 5 million bone marrow cells per recipient were used for secondary or tertiary transplants.

### LDH activity measurement

Bone marrow cells were treated with ACK buffer to lyse erythrocytes and washed with ice-cold 1x PBS prior to LDH activity measurement. LDH activity was measured with a fluorometric LDH activity assay kit according to manufacturer instructions (LSBio, LS-K890-500). 1 million bone marrow cells were homogenized with ice-cold LDH assay buffer and kept on ice for 10 minutes. Samples were centrifuged at 10,000 g for 5 min and the supernatant collected and diluted 10 times. 50 μl of diluted supernatant were mixed with 50 μl of LDH assay reaction mix containing buffer, probe, and substrates. The fluorescence from sample wells or wells containing NADH standard for calibration was measured every 2 minutes for 30 minutes using Spectramax iD3 (Molecular Devices).

### Oxygen consumption measurements

O_2_ consumption measurements were performed with Seahorse XFe96 instrument (Agilent Technologies) according to the manufacturer’s instructions with some modifications. Briefly, a 96 well Seahorse culture plate was coated with poly-L-lysine (Sigma P4707) the day before the analysis. 400,000 Kit^+^ HSPCs or CD45^+^ bone marrow hematopoietic cells were sorted and washed twice with the media used for plating. Cells were added to the culture plate in DMEM (Sigma, D5030) supplemented with 10 mM glucose, 2 mM glutamine and 1 mM pyruvate and the pH was adjusted to 7.4 with 1 M NaOH. Measurements were taken at 37°C every 3 minutes. Respiratory O_2_ consumption was calculated as the drop in O_2_ consumption after addition of 2 μM oligomycin.

### Metabolite consumption and production analysis

500,000 Lin^-^Kit^+^ or CD45^+^ cells were sorted (BD FACSSymphony S6), centrifuged and washed twice with 1 mL cell culture media. For experiments with *Ldha^Δ/Δ^;Ldhb^Δ/Δ^*cells or for *ex vivo* tracing of *Ldha^Δ/Δ^;Ldhb^Δ/Δ^;Pdha1^τι^* cells, DMEM without glucose and pyruvate (Gibco, 11966-025), supplemented with 0.5 mM glucose or U^13^C-glucose was used. For metabolite production analysis of *Ldha^Δ/Δ^;Ldhb^Δ/Δ^;Pdha1^τι^* cells, DMEM (Sigma D5030) supplemented with 2 mM glutamine and 0.2 mM glucose, pH = 7.4 was used. For metabolite production analysis of wild type HSPCs supplemented with lactate and palmitate, DMEM without glucose and pyruvate (Gibco, 11966-025), supplemented with 1 mM glucose, with or without 200 μM palmitate and 5 mM lactate was used. Cells were incubated in 100 μL media + 10 ng/mL SCF + 100 ng/mL TPO in a round bottom 96-well plate at 37°C in a 5% CO_2_ cell culture incubator for 12 hours. Wells containing media without cells were used as a comparison to measure metabolite consumption or production. Media were collected by transferring the samples from each well to individual 1.5 mL tubes and centrifuging cells at 4°C, 400 g. The supernatant was collected into new tubes and centrifuged again. 70 μL of supernatant were collected into new 1.5 ml tubes, frozen immediately in liquid nitrogen and stored at −80°C until LC-MS analysis. For MS analysis, 10 μl of the media were added to a 1.5 mL tube, followed by addition of 90 μL of 90% acetonitrile (Optima LC/MS Water W6-4, Acetonitrile A955-4, Fisher Chemical). An internal standard mix containing stable isotope-labeled glucose (100 μM), pyruvate (20 μM), lactate (20 μM), glutamine (100 μM), glutamate (10 μM), α-KG (1 μM), succinate (1 μM), malate (1 μM), and aspartate (10 μM), was spiked in (2 μL / sample). The samples were vortexed for 30 sec and incubated on ice for 10 minutes, followed by centrifugation at 4°C, 21,000 g for 15 minutes. Supernatants were transferred to new 1.5 ml tubes, and 30 μL were added to an LC vial with an insert for LC-MS analysis. The injection volume was 15 μL.

### BM-ISF and BM collection for metabolomics

BM-ISF was collected from one femur by flushing with 200 μL of cold DPBS followed by centrifugation at 4°C, 400 g, for 5 minutes. The supernatant was transferred into a new 1.5 mL tube. The tubes were centrifuged again and the top part (100μL) of the supernatant was transferred to new 1.5mL tubes and immediately frozen in liquid nitrogen. Metabolites were extracted using 80% acetonitrile:20% water, or 40% methanol:40% acetonitrile:20% water followed by centrifugation. To collect bone marrow cells, pelleted cells from BM-ISF collected tubes were resuspended in 400 μL of ACK buffer for red blood cell lysis for 5 minutes at 4°C followed by addition of 1 mL of cold DPBS and centrifugation. The supernatant was aspirated, and the cell pellet was immediately frozen in liquid nitrogen. Samples were stored in a −80°C freezer until LC-MS analysis.

### Cell isolation for rare cell metabolomics

All cell isolation and preparation for rare cell metabolomics were performed quickly at 4°C in the cold room as described previously^16,17,37^. Mice were euthanized by cervical dislocation and bone marrow cells were obtained by crushing femurs, tibias, vertebrae and pelvic bones with an ice-cold mortar and pestle in a 4°C cold room, on ice in cold 2 ml of Ca^2+^/Mg^2+^-free HBSS supplemented with 0.5% bovine serum albumin. Cells were filtered through a 40-μm cell strainer into 50 ml conical tubes. For sorting hematopoietic stem and progenitor cells, anti-Kit paramagnetic beads were added and gently mixed. Cells were left on ice for 10 minutes followed by addition of fluorescent-conjugated antibodies against surface markers for HSPCs as described above. Cells were incubated on ice for 60 minutes with occasional mixing followed by selection on QuadroMACS magnetic separator (Miltenyi Biotec). For isolation of cells for tracing experiments 1x DPBS (Sigma-Aldrich D8537) supplemented with 0.5% bovine serum albumin was used for cell isolation. Cells were sorted with BD FACSSymphony S6 running with sheath fluid 0.5 x PBS^17^ prepared fresh with Optima LC/MS Water (W6-4, Fisher Chemical) and 10x PBS (Sigma-Aldrich D1408). A 70 μm nozzle with a four-way purity mode was used for cell sorting^17^. Before sorting, the flow sorter was cleaned by sequentially running through it at high speed for 10 minutes each Windex and 10% bleach, followed by 15 minutes of Optima LC/MS Water (W6-4, Fisher Chemical). For rare cell metabolomics 10,000 cells from each of HSC+MPP, HPC1 and myeloid progenitors (MyP, Lin^-^Sca-1^-^Kit^+^) populations were directly sorted into 50 μl of 100% acetonitrile, giving an ∼80% final concentration of acetonitrile. 100% acetonitrile solvent was kept on ice and the sorting temperature was maintained at 4°C throughout the sort. Blank controls consistent of tubes with the same acetonitrile extraction solvent containing sheath fluid from a 4-second test sort. Sorted samples were quickly vortexed and placed on dry-ice throughout the duration of the experiment and then stored at −80°C until analysis.

### Stable isotope tracing *in vivo*

Mice were fasted for 6-8 hours before infusions. Mice were placed under anesthesia using 40 μl injection of 25 mg/ml ketamine and a 27-gauge catheter was inserted in the lateral tail vein for infusions. Mice were infused with U^13^C-glucose (CLM-1396, Cambridge Isotope Laboratories), starting with a bolus for 1 minute with 123 μL/min of tracer to fill the infusion line, followed by continuous infusion of 0.008 mg/g body weight per minute for 3 hours (in a volume of 150 μl/hr). 3 hours after the start of infusion, mice were sacrificed, and all bones were collected. For bone marrow and BM-ISF metabolomics, samples were collected as described above. For HSPC stable isotope tracing, bone marrow cells were obtained by crushing femurs, tibias, vertebrae and pelvic bones with an ice-cold mortar and pestle in a 4°C cold room on ice in cold DPBS supplemented with 0.5% bovine serum albumin. Kit microbeads were used for enrichment followed by staining with surface antibodies. Cells were incubated on ice for 60 minutes with occasional mixing followed by selection on QuadroMACS magnetic separator (Miltenyi Biotec) using DPBS for washing. 10,000-15,000 HSC+MPP or HPC1 cells were sorted into 50 μL of 100% acetonitrile, vortexed and stored in −80^0^C. For myeloid progenitors (MyP, Lin^-^Sca-1^-^Kit^+^) 50,000-100,000 cells were sorted into 100 μL of DPBS, centrifuged immediately after sorting, most supernatant except for ∼20 μl was removed, and metabolites were extracted by addition of 80 μL 100% ACN.

### Metabolite extraction for LC-MS analysis

The metabolite extraction process was performed in a HEPA filtered PCR hood. For HSPC metabolomics, sorted samples were thawed on ice and vortexed for 1 minute. Samples were kept on ice for 10 minutes and then centrifuged at 21,000g for 15 minutes at 4°C. Supernatant was transferred into LC-MS autosampler vials and 20 μL of the extract was injected for LC-MS analysis. Negative control blank samples were processed in parallel with experimental samples. Blank samples contained 0.5 x PBS sheath fluid from a test sort + acetonitrile extraction solvent at equal volumes as the experimental samples. For metabolomics of bone marrow cells, 100 μL of 80% acetonitrile was added to the cell pellet immediately after removing it from the −80°C freezer. Metabolites were extracted with 3 freeze-thaw-vortex cycles (freezing in liquid nitrogen, thawing in ice water), followed by centrifugation at 21,000 g at 4°C for 15 minutes. Supernatants were transferred into a new set of 1.5 mL tubes and centrifuged again and 25 μL were transferred into LC vials containing inserts on ice. 15 μL were injected into the LC-MS. Corresponding sample blank for this experiment was prepared by adding the same amount of 80% acetonitrile into an empty 1.5 mL tube processed side by side with the samples. For BM-ISF extraction, 10 μL of BM-ISF was added to 90 μL of 90% acetonitrile. After vortexing for 30s, the samples were kept on ice for 10 minutes before centrifuging at 21,000 g at 4°C for 15 minutes. The supernatant was transferred into a new set of 1.5 mL tubes, and 25 μL were transferred to LC-MS vials. A sample blank was prepared by mixing 10 μL DPBS with 90 μL of 90% Acetonitrile and processed the same way as the BM-ISF samples. 15 μL were injected for LC-MS analysis. For serum extraction, 67 μL of 80% acetonitrile was added to 3 μL of serum, vortexed for 30s, left on ice for 10 minutes, and centrifuged at 21,000 g at 4°C for 15 minutes. The supernatant was transferred into a new set of 1.5 mL tubes, centrifuged again, and 30 μL were transferred to LC-MS vials. 10 μL were injected for LC-MS analysis. A sample blank was prepared by mixing 3 μL HPLC-grade water with 67 μL of 80% Acetonitrile and processed the same way as the serum samples.

### LC-MS analysis

Metabolites were separated prior to mass spectrometry using a Thermo Scientific (Bremen, Germany) Vanquish Flex liquid chromatography (LC) system. LC was performed on a Millipore ZIC-pHILIC column (5 μm, 2.1×150 mm), a binary solvent system of 10 mM ammonium acetate in water, pH 9.8 (solvent A) and acetonitrile (solvent B) with a constant flow rate of 0.25 mL/min or 0.4 mL/min was used. The column was equilibrated with 90% solvent B. The liquid chromatography gradient was: 0–15 minutes linear ramp from 90% B to 30% B; 15–18 minutes isocratic flow of 30% B; 18–19 minutes linear ramp from 30% B to 90% B; 19–27 column regeneration with isocratic flow of 90% B. Metabolites were analyzed on a Thermo Scientific Orbitrap Exploris 480 mass spectrometer or QExactive HF-X, collecting both positive and negative ion spectra, as previously described^16,52^.

### Metabolomics data processing

TraceFinder 5.1 (Thermo) equipped with an in-house spectral library was used to identify and quantify metabolites. The library compares characterized precursor and product ion spectra to spectra obtained from chemical standards and biological extracts. All metabolites were identified from raw data files with a 5 ppm mass tolerance. Raw values for metabolite MS peak abundance (area under the curve) were first cleaned by filtering out metabolites for which signal intensity was lower than 2 x the signal intensity in the blank samples. For nutrient consumption experiments, metabolites for which signal intensity was lower than 3 x the signal intensity in the blank samples were filtered out. Signal intensities were normalized to the median signal intensity to account for variability in total sample amount. For *ex vivo* metabolite production data, metabolites were defined as net produced if their average signals were higher than those of the media-only controls. Labeled internal standards were used for absolute quantitation. For rare cell metabolomics, raw data were first cleaned by filtering out metabolites for which signal intensity was lower than 2 x the signal intensity in the blank sample, and then polar metabolites and non-polar lipids were normalized separately using the corresponding median. Metaboanalyst 6.0 was used to process metabolomics data, including missing value imputation (features with missing values in more than 50% of the samples were removed, and for the remaining missing values, a value equal to 0.2 of the minimum value for that feature within the treatment group was used), log transformation, and autoscaling for display. For stable isotope tracing data, natural abundance was corrected by using Accucor^53^. In graphs depicting total metabolite quantities of each isotopologue from tracing experiments, isotopologues for each metabolite were added and normalization using the median total signal intensity of quantified metabolites was used to correct for variability in sample quantity.

### Protein Extraction and Western Blot Analysis

Bone marrow cells were lysed using ACK lysis buffer and equal number of cells were collected and washed with ice-cold 1x PBS. Cells were pelleted and extraction was done with RIPA buffer containing phosphatase and protease inhibitors. Samples were vortexed at 10 minutes interval on ice for 30 minutes and centrifuged at 17,000g for 15 minutes. Supernatant was collected and PLB loading buffer (BioRad) and β-mercaptoethanol were added followed by heating samples at 70°C for 10 minutes. Samples were run on 4-15% Mini-Protean TGX gels (BioRad) for separated and transferred to 0.2 mm PVDF membranes (BioRad) by wet transfer using Tris Glycine transfer buffer (BioRad). Western blots were performed using antibodies against LDHA (Fisher 50-173-3595), LDHΑ/B (Santa Cruz Sc-100775), PDH-E1α (Abcam, ab110330), Vinculin (CST, 4650S). Signals were detected using SuperSignal West Pico or SuperSignal West Femto chemiluminescence kits (Thermo Scientific).

### AZD3865 treatment

Mice were administered 30 mg/kg of AZD3965 (Selleckchem, S7339) by oral gavage for 7 continuous days. AZD3965 was dissolved in 5% dimethyl sulfoxide (DMSO) (Sigma, D2650) + 40% PEG300 (Selleckchem, S6704) + 5% Tween 80 (Selleckchem, S6702) + 50% molecular biology grade water (Cytiva, SH30538.02).

### PCR for genotyping

The following primers were used for genotyping: *Ldha*: 5’-CTGAGCACACCCATGTGAGA-3’, 5’-AGCAACACTCCAAGTCAGGA-3’, 5’-CAGAGAGGGATGCCTTTCTCATAC-3’. Wild-type band=458 bp, Flox band=648 bp, Knockout band=796 bp. *Ldhb*: 5’-CTGAGCTGCTCAGAGCCAAC-3’, 5’-GATGGCTTAGCCCCAATCTACC-3’, 5’-GGACCATTTCTAAGTAGATGGAAATTCTG-3’. Wild-type band=140 bp, Flox band=325 bp, Knockout band=470 bp. *Pdha1*: 5’-ACAGATTACACCCGACTGCC-3’, 5’-GCTGGCTCACCTGGTAAACA-3’, 5’-GGTCCCGAGAAGTGAGCAAA-3’. Wildtype band=205 bp, Flox band=290 bp, Knockout band=523 bp. *Slc16a1* (MCT1): 5’-GCA GCA TGT GGT CCT CTC TTA AG-3’, 5’-GTC CTC ACC TCT CTG TGC-3’, 5’-TGG TTC TCT TGT TAT CAG TGT TGG GTG-3’. Wildtype band=∼250 bp, Flox band=∼350 bp, Knockout band=∼320 bp.

### Statistical analysis

To assess the significance of a difference in means between treatments, a t-test or 1-way ANOVA was used when the data did not significantly deviate from normality and did not have significantly unequal variances, a Welch’s t-test or Brown–Forsythe ANOVA when the data had unequal variances, and a Mann–Whitney or Kruskal–Wallis test when the data significantly deviated from normality. To test if data deviated from normality (p<0.01 for at least one treatment), we used the D’Agostino-Pearson test or, when n < 8, the Shapiro–Wilk test. To test if variance significantly differed among treatments, we used the F-test (for experiments with two treatments) or the Brown–Forsythe test (for more than two treatments). In metabolomics analysis, multiple comparisons correction was performed by controlling the false discovery rate at 5% using the method of Benjamini, Krieger, and Yekutieli. Graphs show *p < 0.05, **p < 0.01, ***p < 0.001 unless noted otherwise. All statistical analysis was performed with measurements from biological replicates. All statistical tests comparing two populations were two-sided. Statistical analyses were performed with Graphpad Prism v10 unless noted otherwise. Graphs were plotted using Prism.

## Acknowledgements.

The work was funded by grants from the Cancer Prevention and Research Institute of Texas (CPRIT) (Scholar Award RR180007), American Society of Hematology (Faculty Scholar award), Moody Foundation, Welch Foundation (I-2053-20210327), Alex’s Lemonade Stand Foundation (‘A’ Award), the Haggerty Foundation, the Rally Foundation (25IN22), and the National Institutes of Health (R01DK125713, R01HL161387) to MA; an American Society of Hematology Inclusion Pathway (HIP) Fellow award to EOK; and an American Society of Hematology Fellow to Faculty Scholar award to YL. We thank Tom Mathews and the Children’s Research Institute (CRI) Metabolomics facility (supported by the CPRIT Core Facilities Support Award RP240494) for mass spectrometry support, Michael Ortiz and the CRI Flow cytometry facility for flow cytometry support, Landon Nguyen, Joseph Rose III, Jacob Zielke, and Grace Ding for mouse colony management and technical assistance, Brett Morrison (Johns Hopkins) for sharing *Slc16a1^fl^* mice, and Sean Morrison and Prashant Mishra for comments on the manuscript.

---

**Figure S1.**
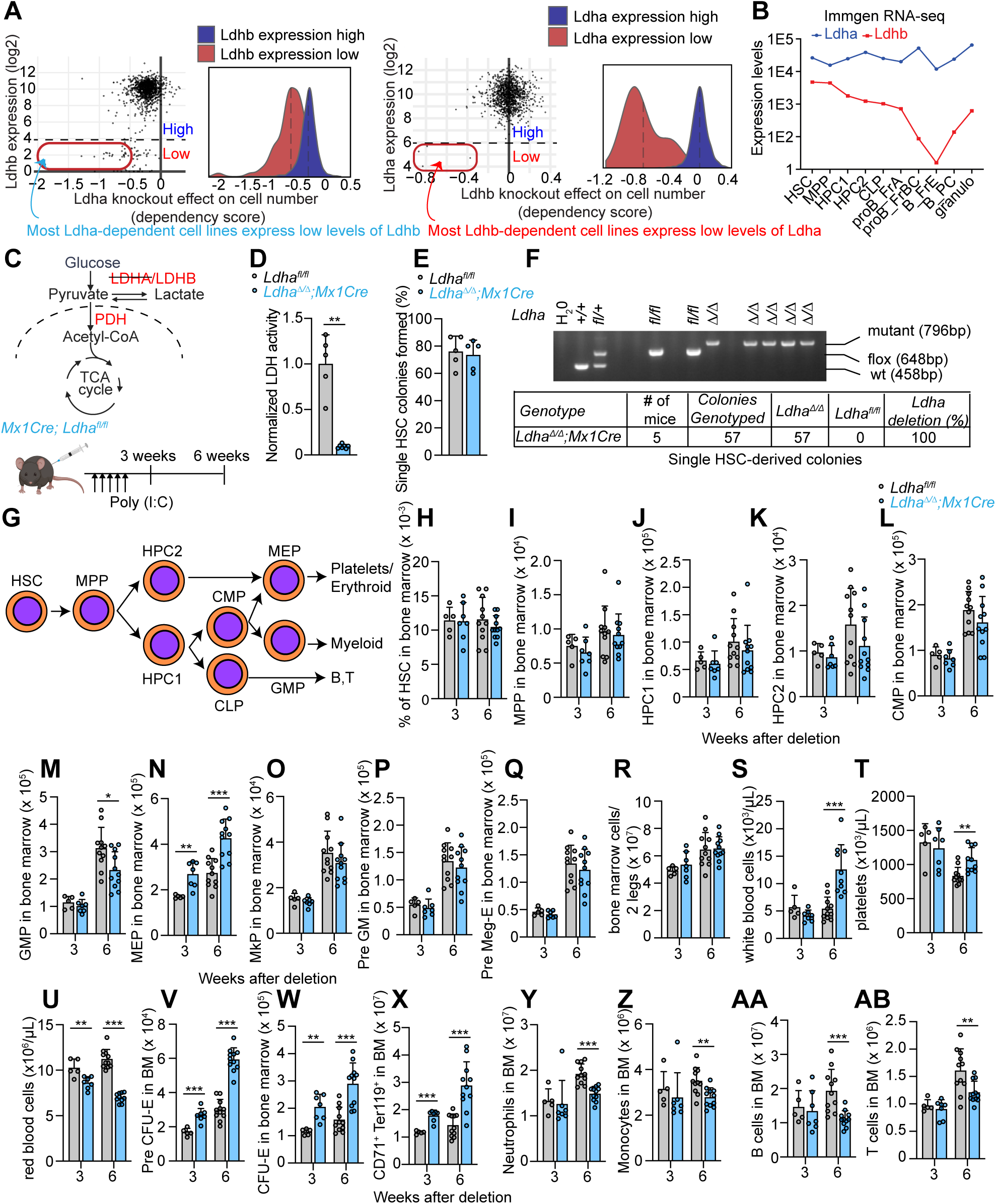
The effects of conditional *Ldha* deletion on bone marrow hematopoiesis. **A,** DepMap analysis showing that *in vitro* most cell lines dependent on LDHA have low LDHB expression, and vice versa. **B,** Expression profiles of Ldha and Ldhb in mouse hematopoietic cells in the Immgen database **C,** Schematic of glycolysis termination illustrating conditional *Ldha* deletion with *Mx1Cre*. **D,** LDH activity assay showing that *Ldha* deletion sharply reduces total LDH enzymatic activity of bone marrow hematopoietic cells following ACK lysis. **E,** Colony-forming capacity in methylcellullose of single sorted *Ldha^Δ/Δ^* HSCs. **F,** Genotyping of colonies from **E** confirming 100% deletion in HSCs 3 weeks after deletion. **G,** Lineage relationships of analyzed hematopoietic cell types. **H,** Frequency of HSCs in the bone marrow at 3 and 6 weeks post deletion. **H-Q,** Numbers of multipotent and restricted progenitors in the bone marrow at 3 and 6 weeks post *Ldha* deletion. Stem and progenitor cells are immunophenotypically defined as: CD150^+^CD48^-^Lineage^-^Sca-1^+^Kit^+^ HSCs, CD150^-^CD48^-^Lineage^-^Sca-1^+^Kit^+^ multipotent progenitors (MPPs), CD150^-^CD48^+^Lineage^-^Sca-1^+^Kit^+^ oligopotent hematopoietic progenitors-1 (HPC1), CD150^+^CD48^+^Lineage^-^Sca-1^+^Kit^+^ oligopotent hematopoietic progenitors-2 (HPC2). See methods for immunophenotypic definitions of restricted progenitors. **R-U,** bone marrow cellularity and peripheral blood cell counts. **V-X** bone marrow erythroid progenitor cells. **Y-AB,** bone marrow neutrophils, monocytes, B and T cells. Data represent mean ± s.d. Statistical significance was assessed with Welch’s test (**D**, **X**), or t-test (all others). Significance is indicated as *p < 0.05, **p < 0.01, ***p < 0.001.

**Figure S2.**
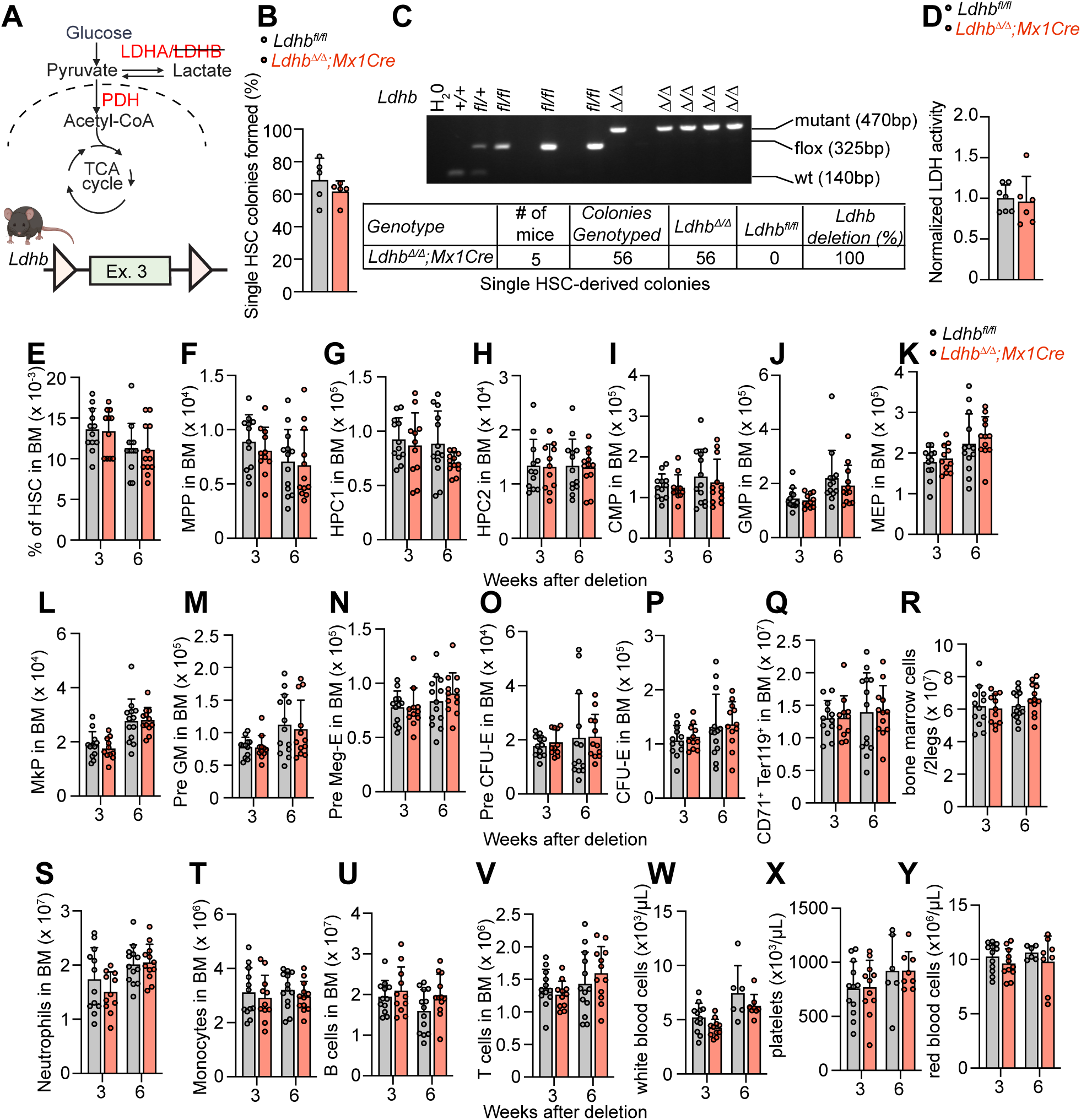
The effects of conditional *Ldhb* deletion on bone marrow hematopoiesis. **A,** Schematic of glycolysis termination illustrating conditional *Ldhb* deletion with *Mx1Cre*. **B,** Colony-forming capacity in methylcellullose of single sorted *Ldhb^Δ/Δ^*HSCs. **D,** LDH activity assay shows total LDH activity is unchanged in *Mx1Cre;Ldhb^Δ/Δ^* bone marrow hematopoietic cells. **E,** Frequency of HSCs in the bone marrow at 3 and 6 weeks post deletion. **F-Q,** Numbers of multipotent and restricted progenitors in the bone marrow of *Mx1Cre;Ldhb^Δ/Δ^* mice. **R–V,** bone marrow cellularity and numbers of neutrophils, monocytes, B, and T cells in *Mx1Cre;Ldhb^Δ/Δ^* mice. **X–Y,** peripheral blood cell counts in *Mx1Cre;Ldhb^Δ/Δ^* mice. Data represent mean ± s.d.

**Figure S3.**
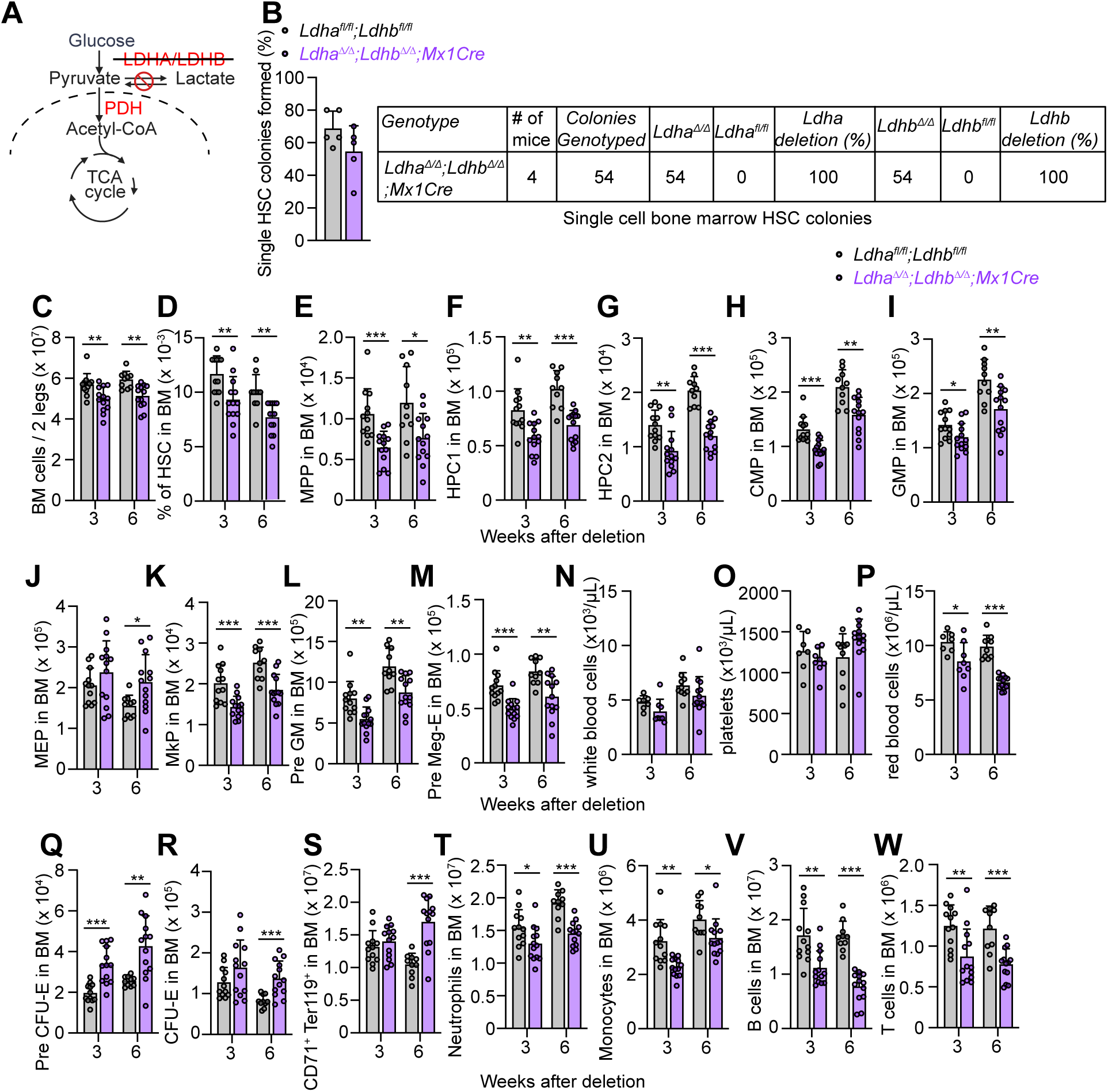
The effects of conditional *Ldha;Ldhb* deletion on bone marrow hematopoiesis. **A,** Schematic of glycolysis termination illustrating conditional *Ldha;Ldhb* deletion with *Mx1Cre*. **B,** Colony forming capacity of single HSCs sorted in methylcellulose after *Ldha* and *Ldhb* deletion, and colony genotyping confirming 100% deletion in *Mx1Cre;Ldha^Δ/Δ^;Ldhb^Δ/Δ^* HSCs. **C-M,** Analysis of bone marrow cellularity and HSC and progenitor cell types in the bone marrow of *Mx1Cre;Ldha^Δ/Δ^;Ldhb^Δ/Δ^* mice 3 and 6 weeks after deletion. **N-P,** Peripheral cell counts. **Q-S,** Numbers of erythroid progenitors. **T-W,** Numbers of neutrophils, monocytes, B, and T cells in *Mx1Cre;Ldhb^Δ/Δ^* mice. All data represent mean ± s.d. Statistical significance was assessed with a Welch’s test (**Q-R**) or a t-test (all others). All figures show * p<0.05, ** p<0.01, ***p<0.001.

**Figure S4.**
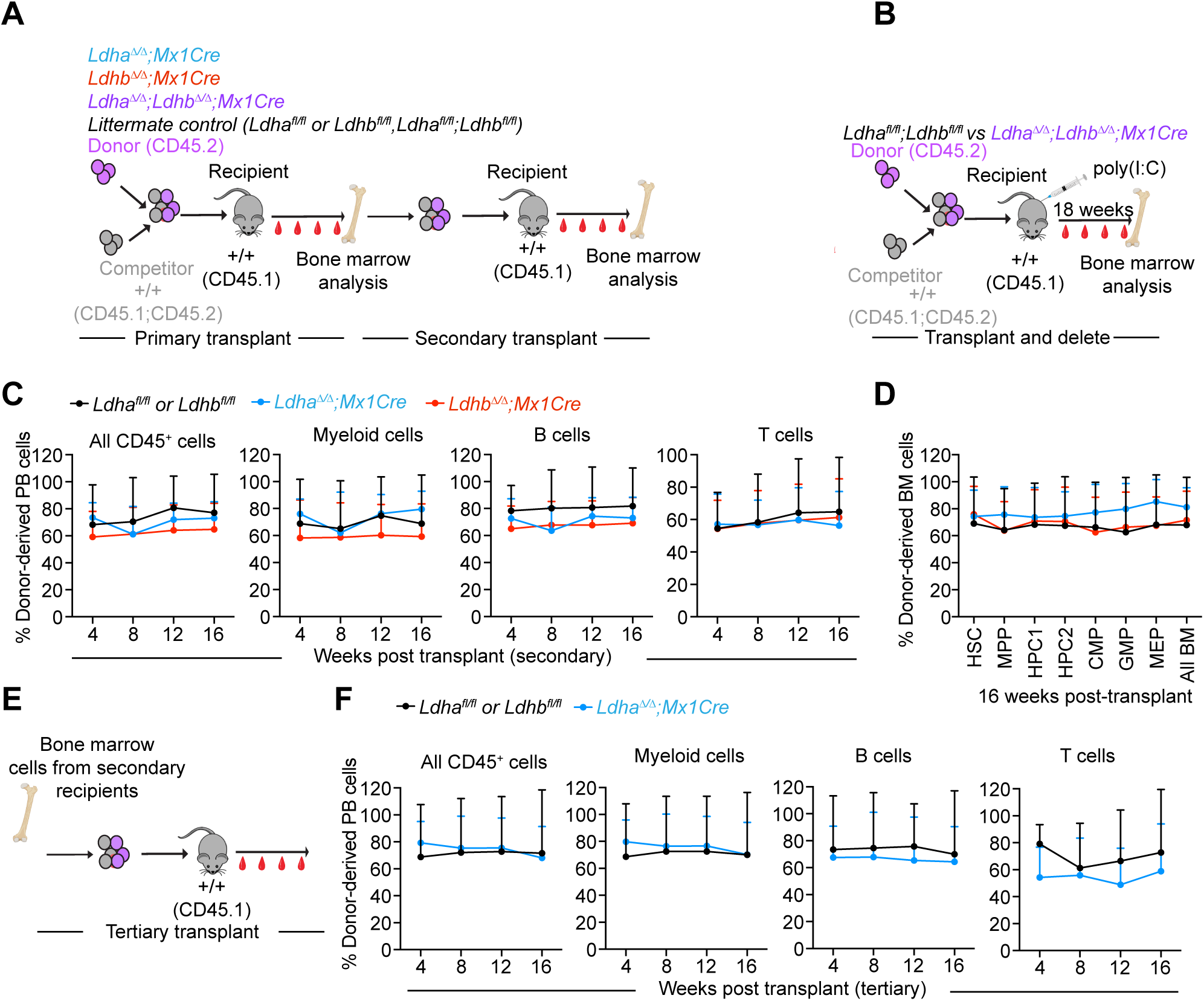
*Ldha* and *Ldhb* are dispensable for HSC reconstitution in serial transplants. **A,** Experimental design for primary and secondary transplant assays in which the noted genes are deleted before transplants. **B,** Experimental design for primary transplant assays in which the noted genes are deleted 6 weeks after transplants in chimeric transplant recipients. and HSPC chimerism in the bone marrow **(D)**. (n = 7-20 mice/genotype for blood analysis and 7-15 mice/genotype for bone marrow analysis from 3 independent experiments) **C–D,** Secondary transplant: donor-derived CD45+, myeloid, B, and T cell chimerism in blood **(C)** and HSPC chimerism in the bone marrow **(D)**. (n = 7-20 mice/genotype for blood analysis and 7-15 mice/genotype for bone marrow analysis from 3 independent experiments **E,** Experimental design for tertiary transplants. **F,** Tertiary transplant for *Ldha* deletion: Blood chimerism over 16 weeks post-transplant. (n=5-9 mice/genotype for blood analysis from 3 independent experiments). All data represent mean ± s.d. Statistical significance was assessed with a one-way ANOVA test (**C-D**) or t-tests (**F**). All figures show * p<0.05, ** p<0.01, ***p<0.001.

**Figure S5.**
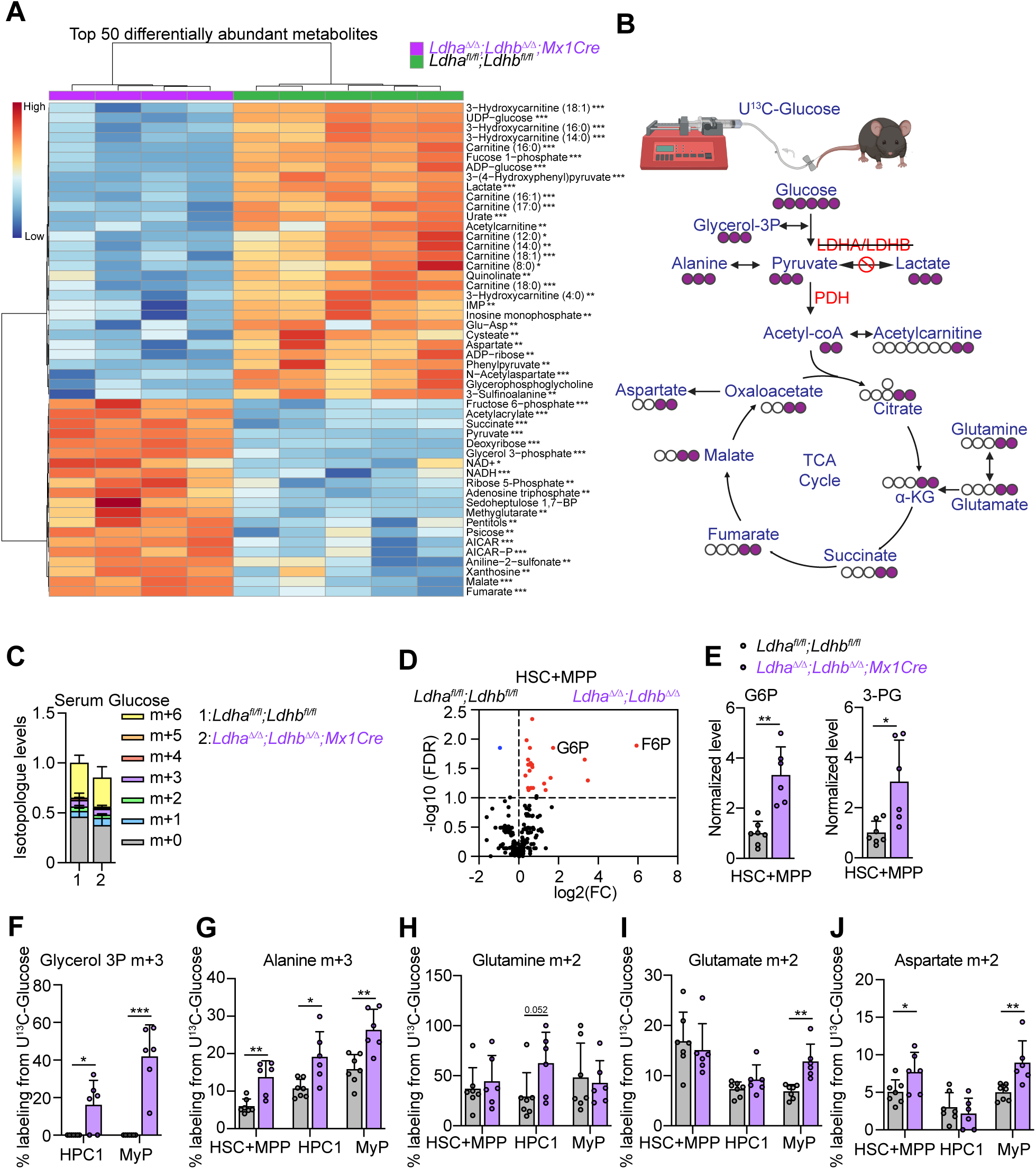
Metabolomics and U13C-glucose tracing analysis in *Mx1Cre; Ldha^Δ/Δ^;Ldhb^Δ/Δ^*HSCs, restricted progenitors and total bone marrow cells. **A,** Heatmap of metabolomics analysis showing the top 50 significantly altered metabolites in *Ldha;Ldhb-*deficient bone marrow cells. **B,** U^13^C-glucose *in vivo* tracing showing the flow of labeled carbons in the indicated metabolites. **C,** No changes in serum glucose labeling in U^13^C-glucose *in vivo* tracing. **D-E,** Metabolomics analysis of *Ldha^Δ/Δ^;Ldhb^Δ/Δ^* HSCs+MPPs. Volcano plot shows fold change in *Ldha^Δ/Δ^;Ldhb^Δ/Δ^* vs control HSC+MPPs. **F-J,** Labeling in indicated metabolites in HSCs and progenitors after U^13^C-glucose *in vivo* tracing (n = 6-7 mice/genotype). Data represent mean ± s.d.; Statistical significance was assessed with multiple t-tests followed by FDR correction at 5% with the method of Benjamini, Krieger, and Yekutieli (**A**, **D**) or Welch’s test (**E**) or a Mann Whitney test (**F**) or a t-test (rest). Data represent mean ± s.d.

**Figure S6.**
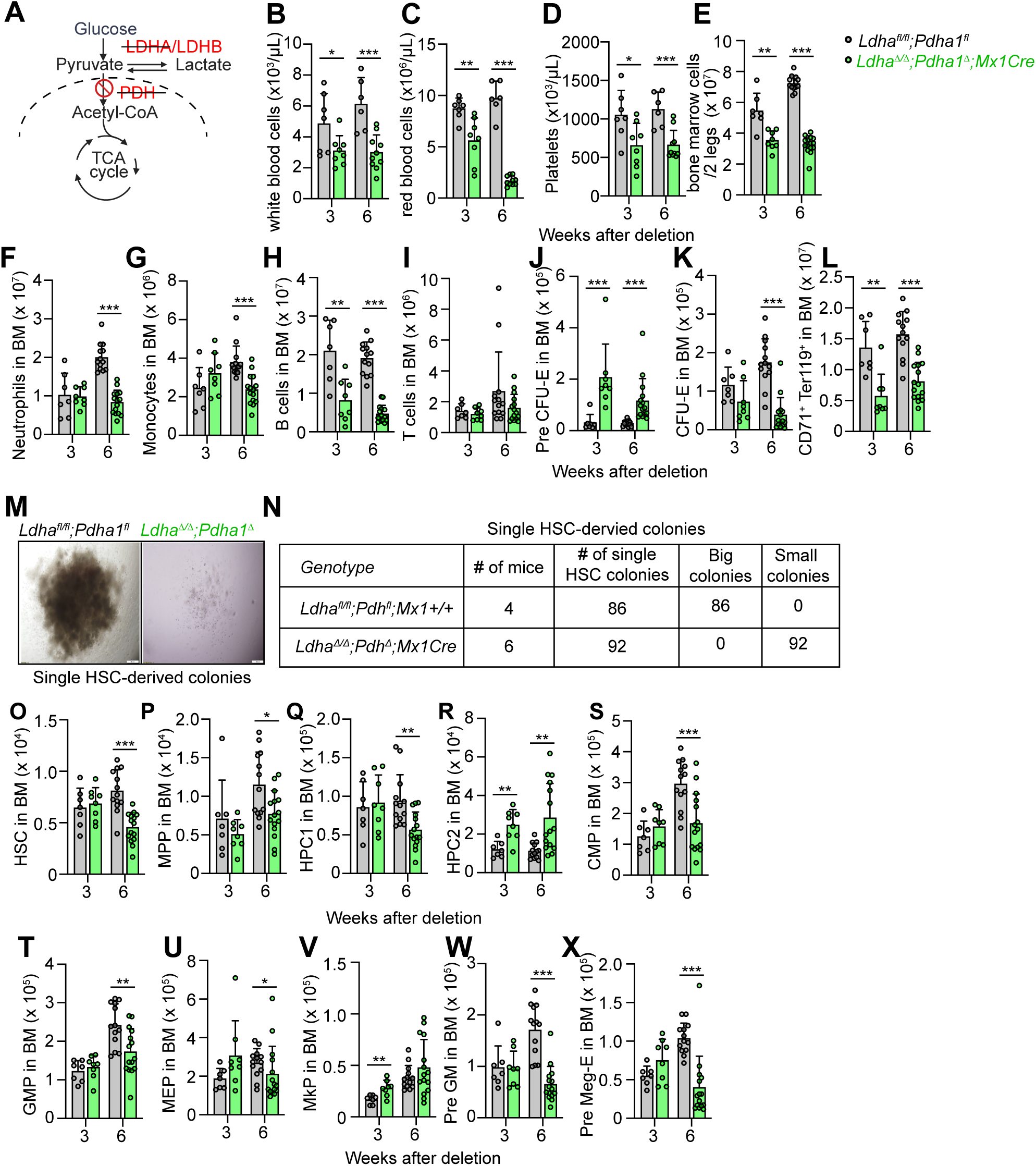
The effects of conditional *Ldha;Pdha1* deletion on bone marrow hematopoiesis. **A,** Schematic of glycolysis termination illustrating LDHA and PDH deletion with *Mx1Cre;Ldha^Δ/Δ^;Pdha1^Δ^* mice. **B-D,** Peripheral blood counts. **E,** Bone marrow cellularity. **F–I,** Numbers of neutrophils, monocytes, B, and T cells. **J–L,** Numbers of bone marrow erythroid progenitors. **M-N,** Images and quantification of methylcellulose colonies from single sorted *Mx1Cre;Ldha^Δ/Δ^;Pdha1^Δ/Δ^* HSCs or controls. **O-X,** HSC and progenitor number in the bone marrow of *Mx1Cre;Ldha^Δ/Δ^;Pdha1^Δ^* mice. N = 6-10 mice/genotype per time point for b-l and 7-15 mice/genotype for o-x. All data represent mean ± s.d. Statistical significance was assessed with a Welch’s test (**C**) or a t-test (rest).

**Figure S7.**
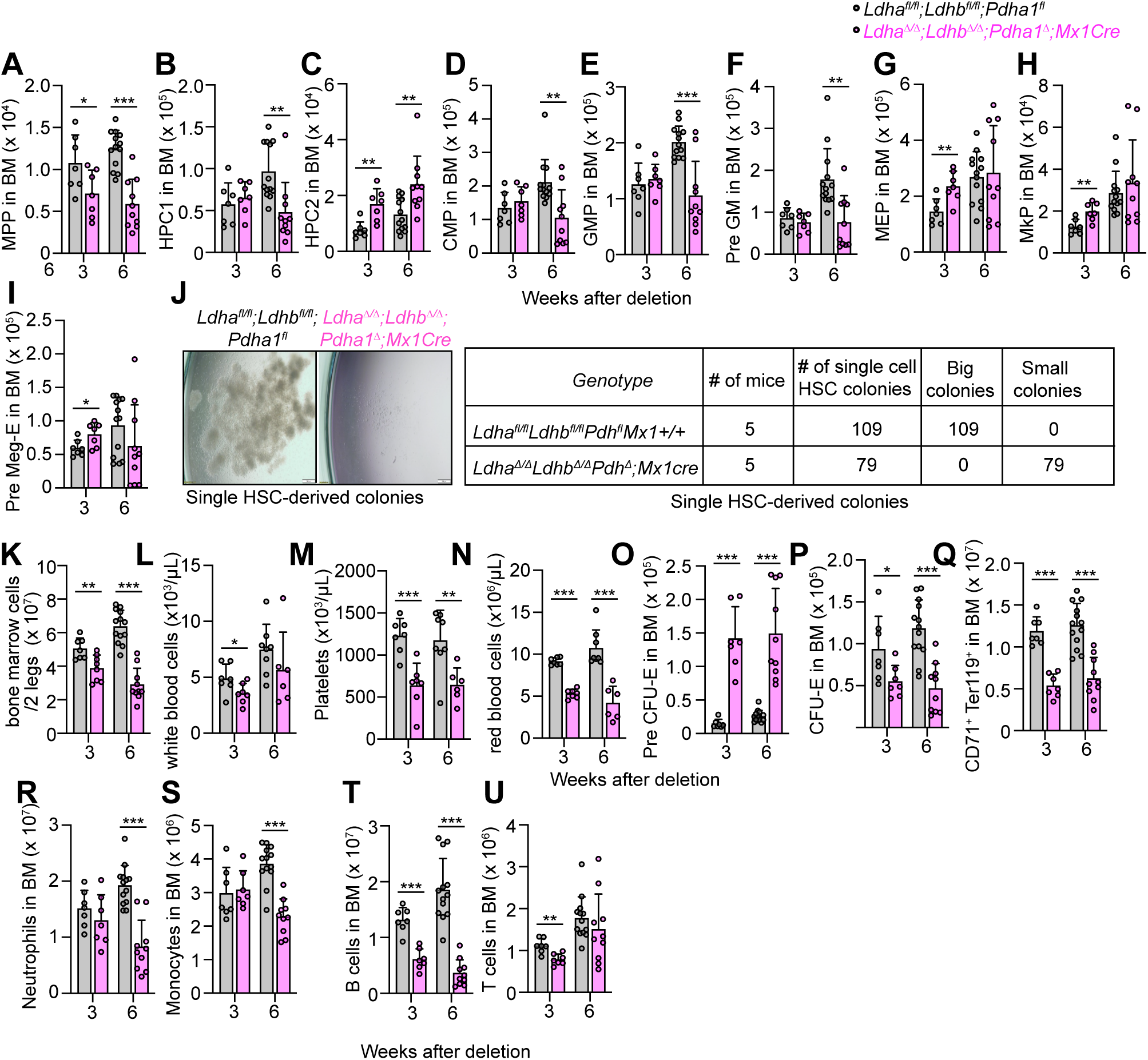
The effects of conditional *Ldha;Ldhb;Pdha1* deletion on bone marrow hematopoiesis. **A–I,** Numbers of multipotent and restricted myeloid progenitors in the bone marrow of *Mx1Cre;Ldha^Δ/Δ^;Ldhb^Δ/Δ^;Pdha1^Δ^* mice at 3 and 6 weeks after deletion. **J,** *Images and quantification of methylcellulose colonies from single sorted Mx1Cre;Ldha^Δ/Δ^*;*Ldhb^Δ/Δ^ ;Pdha1^τι^* HSCs or controls. **K,** Bone marrow cellularity. **L-N,** Peripheral blood counts. **O–Q,** Numbers of erythroid progenitors. **R-U,** Numbers of neutrophils, monocytes, B, and T cells in *Mx1Cre;Ldha^Δ/Δ^;Ldhb^Δ/Δ^;Pdha1^Δ^* mice. N = 7 mice/genotype at 3 weeks and 10-13 mice/genotype at 6 weeks after deletion. All data represent mean ± s.d. Statistical significance was assessed with a Mann-Whitney test (**O**) or a t-test (rest). All figures show * p<0.05, ** p<0.01, ***p<0.001.

**Figure S8.**
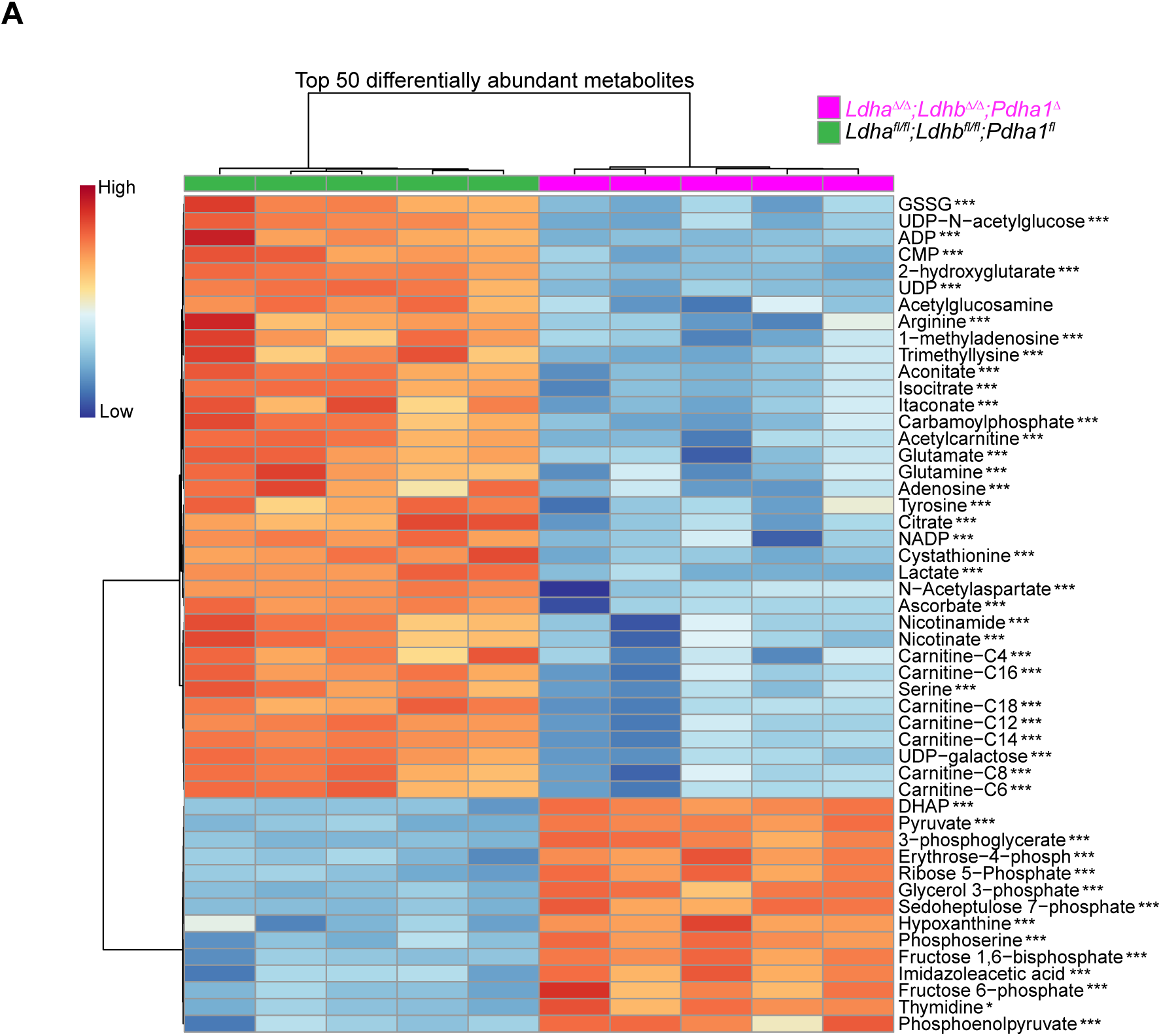
Triple LDHA, LDHB, and PDH deletion reprograms the hematopoietic cell metabolome. Heatmap of metabolomics analysis showing the top 50 significantly altered metabolites in *Ldha;Ldhb;Pdha1-*deficient bone marrow cells. Data represent mean ± s.d.; Statistical significance was assessed with multiple t-tests followed by FDR correction. Significance: *p < 0.05, **p < 0.01, ***p < 0.001.

**Figure S9.**
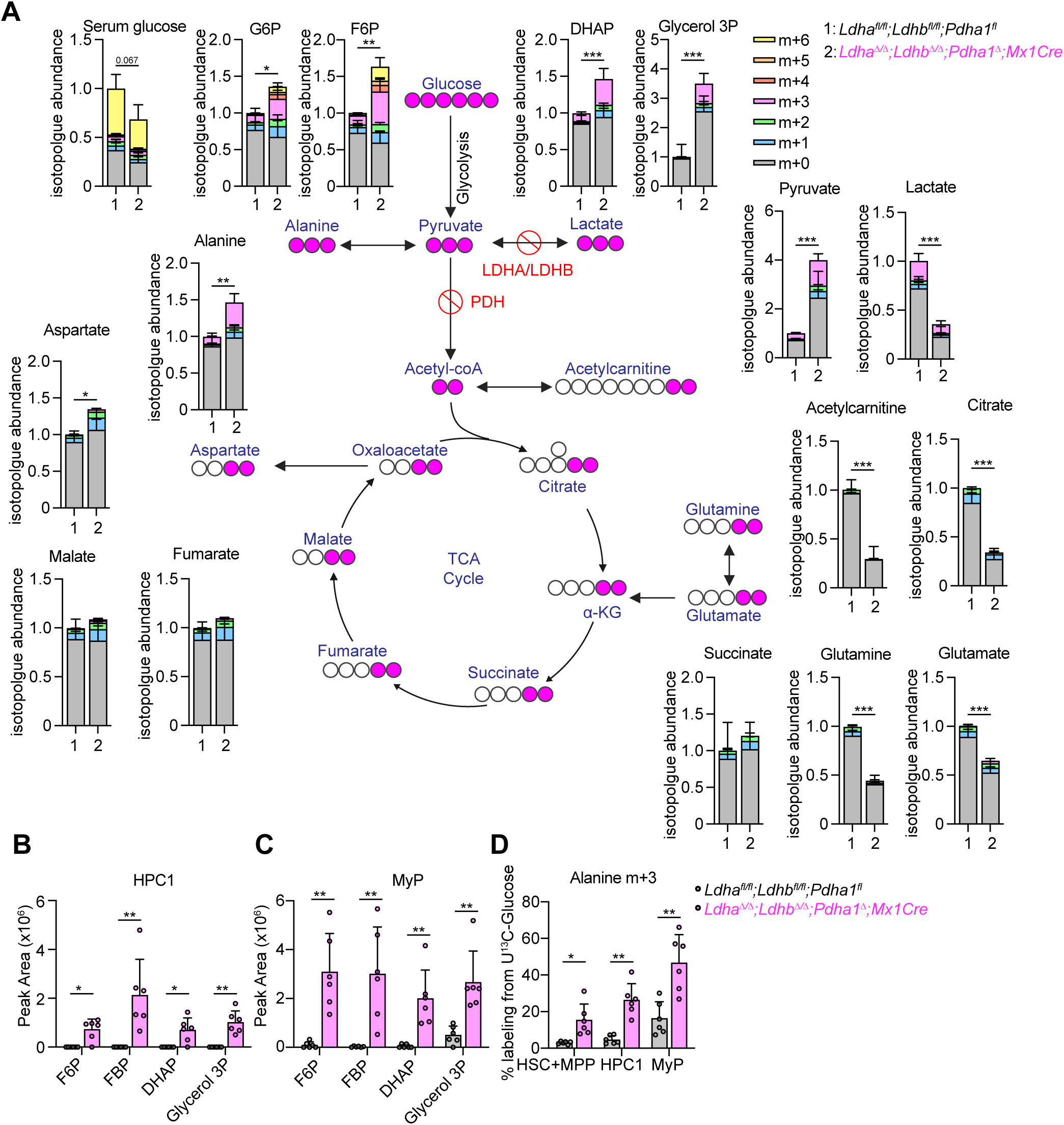
U13C-glucose tracing and metabolomics analysis in hematopoietic cells from *Mx1Cre;Ldha^Δ/Δ^;Ldhb^Δ/Δ^;Pdha1^Δ^* or littermate control mice *in vivo*. **A,** U^13^C-glucose-derived labeling in central carbon metabolites in hematopoietic cells from the bone marrow. Shown are isotopologue abundances of indicated metabolites and serum glucose labeling. **B–C,** Rare-cell metabolomics showing elevated upper glycolytic metabolites in HPC1 (**B**) and myeloid progenitors (MyP) (**C**). **D,** Fractional enrichment of alanine in HSCs and progenitors after U^13^C-glucose tracing *in vivo,* normalized to serum glucose labeling. Data represent mean ± s.d.; Statistical significance was assessed with t-tests (**A**) or a Mann-Whitney test (**B**, **C**) or a Welch’s test (**D**). Significance: *p < 0.05, **p < 0.01, ***p < 0.001.

**Figure. S10.**
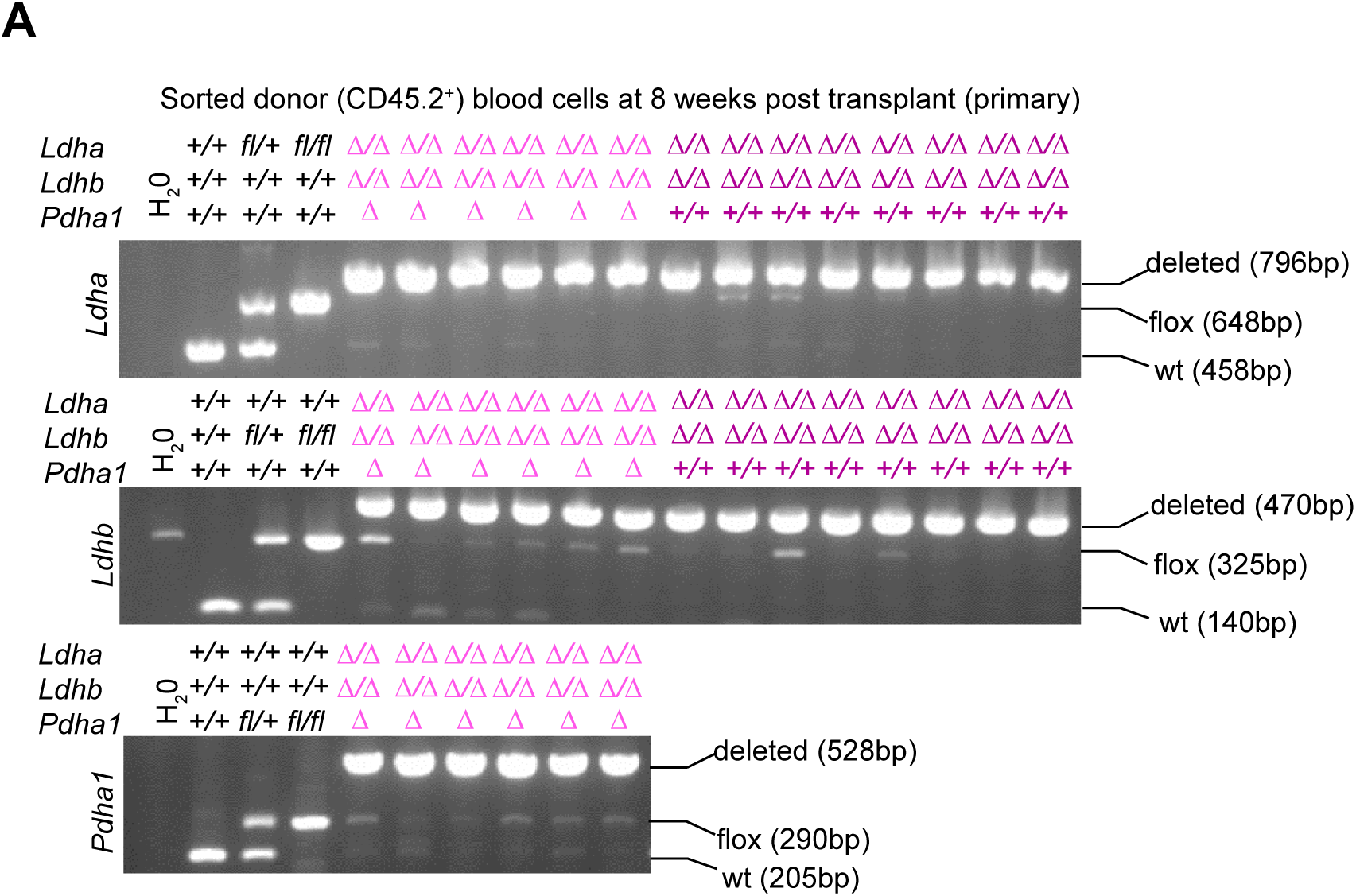
Verification of deletion efficiency in donor-derived blood cells. **A,** 20,000 CD45.2^+^ donor-derived peripheral blood cells from double and triple knockout recipients were sorted 8 weeks post-transplant and genotyped for the indicated alleles. At least 2 of 5 recipient mice per experiment were analyzed (3 independent experiments).

**Figure S11.**
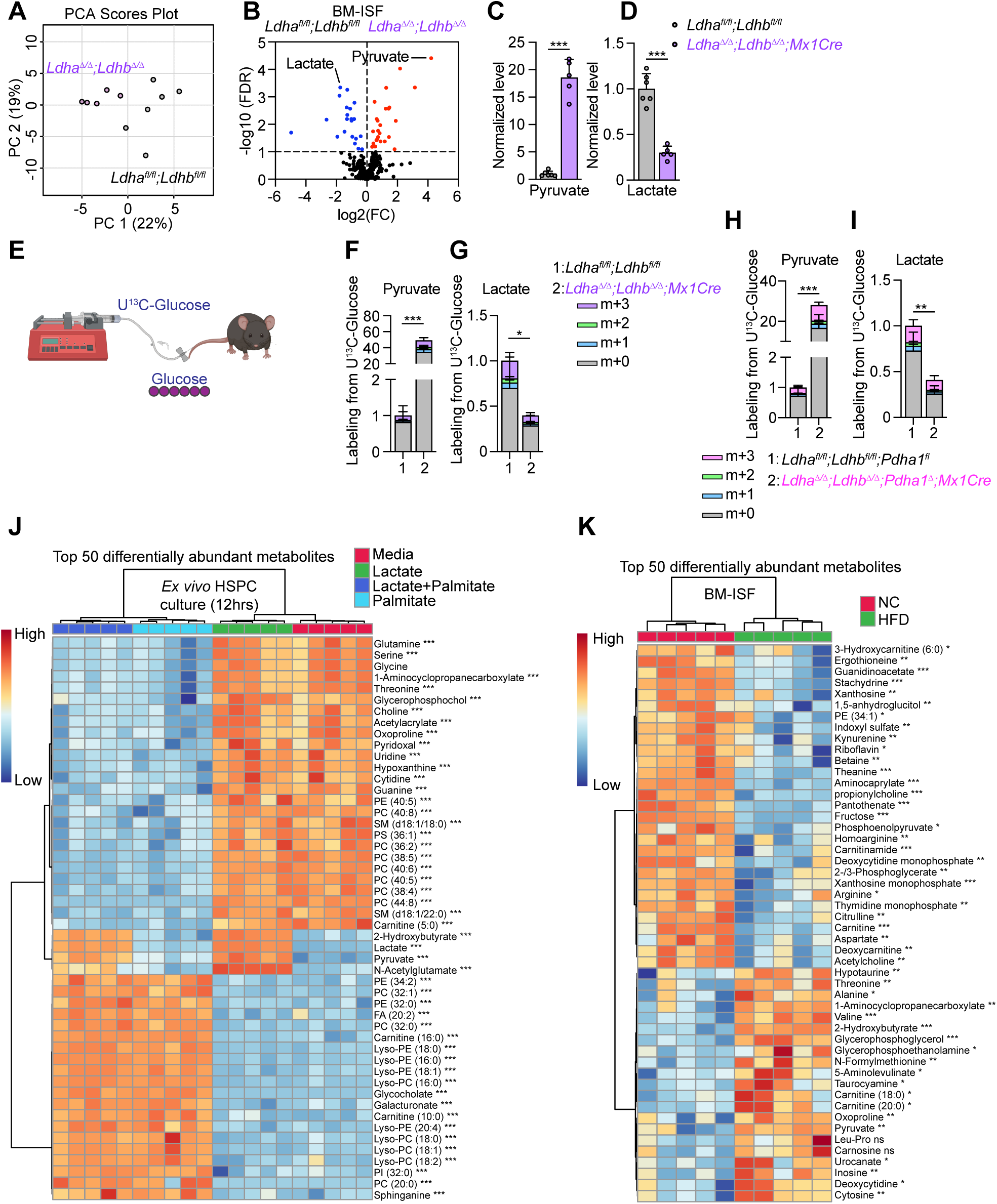
Glucose-derived pyruvate accumulates and glucose-derived lactate is depleted in the *Ldha^Δ/Δ^;Ldhb^Δ/Δ^* and *Ldha^Δ/Δ^;Ldhb^Δ/Δ^;Pdha1^Δ^* bone marrow ISF. **A,** PCA of BM-ISF metabolites in *Mx1Cre;Ldha^Δ/Δ^;Ldhb^Δ/Δ^* mice and littermate controls. **B,** Volcano plot showing pyruvate as the most elevated and lactate as the most depleted metabolite in *Ldha^Δ/Δ^;Ldhb^Δ/Δ^* BM-ISF. **C–D,** BM-ISF levels of pyruvate and lactate in *Ldha^Δ/Δ^;Ldhb^Δ/Δ^* mice. **E–G,** Pyruvate and lactate isotopologue levels in *Ldha^Δ/Δ^;Ldhb^Δ/Δ^*ISF after tracing with U^13^C-glucose. Values were normalized to the sum of the isotopologues in the control. **H–I,** Pyruvate and lactate isotopologue levels in *Ldha^Δ/Δ^;Ldhb^Δ/Δ^;Pdha1^Δ^*ISF after tracing with U^13^C-glucose. Values were normalized to the sum of the isotopologues in the control. **J,** Heatmap of metabolomics analysis showing the top 50 significantly altered metabolites in *ex vivo* culture of sorted Lin^-^Kit^+^ HSPCs bone marrow cells in media supplemented with 1 mM glucose media with or without 200 μM palmitate or 5 mM lactate for 12 hours. **K,** Heatmap of metabolomics analysis showing the top 50 significantly altered metabolites in after 1 week of high fat diet (HFD) or normal chow (NC) feeding. Data represent mean ± s.d.; Statistical significance was assessed with Welch’s test (**C-I**), Tukey’s HSD ANOVA test with FDR correction (**J**) and multiple t-tests followed by FDR correction (**K**). Significance: *p < 0.05, **p < 0.01, ***p < 0.001.

**Figure S12.**
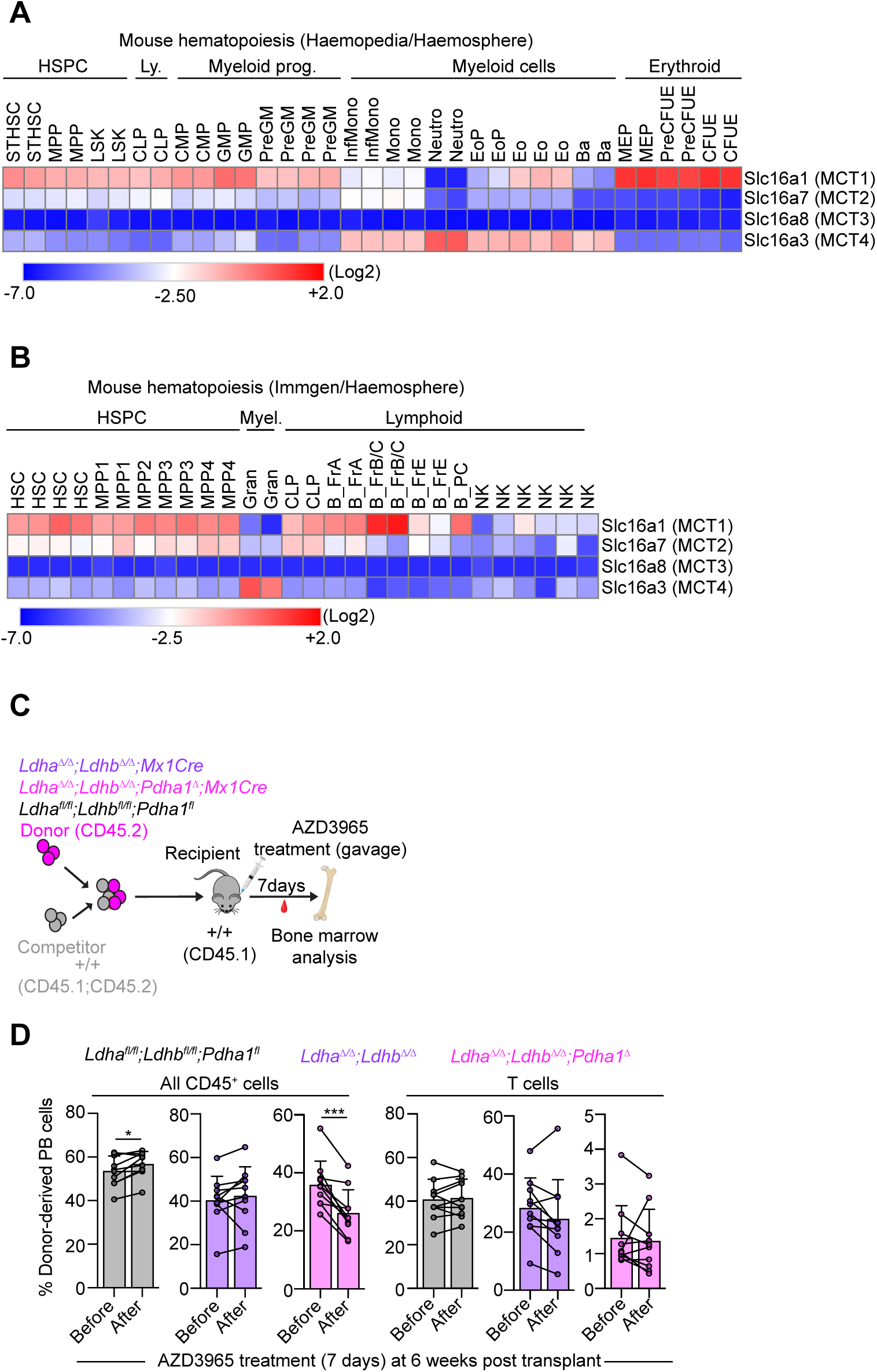
MCT expression in hematopoiesis and effects of AZD3965 treatment on HSC function. **A–B,** Gene expression of MCT1–4 in hematopoietic cell types. Data from the Immgen or the Haemopedia RNA-seq datasets obtained from the Haemosphere database. **C,** Experimental design for MCT1 inhibition with AZD3965 *in vivo*. **D,** Donor-derived peripheral blood CD45^+^ and T cell chimerism before or after AZD3965 treatment. Data represent mean ± s.d.; Statistical significance was assessed with paired t-tests (**D**). Significance: *p < 0.05, **p < 0.01, ***p < 0.001.

**Figure S13.**
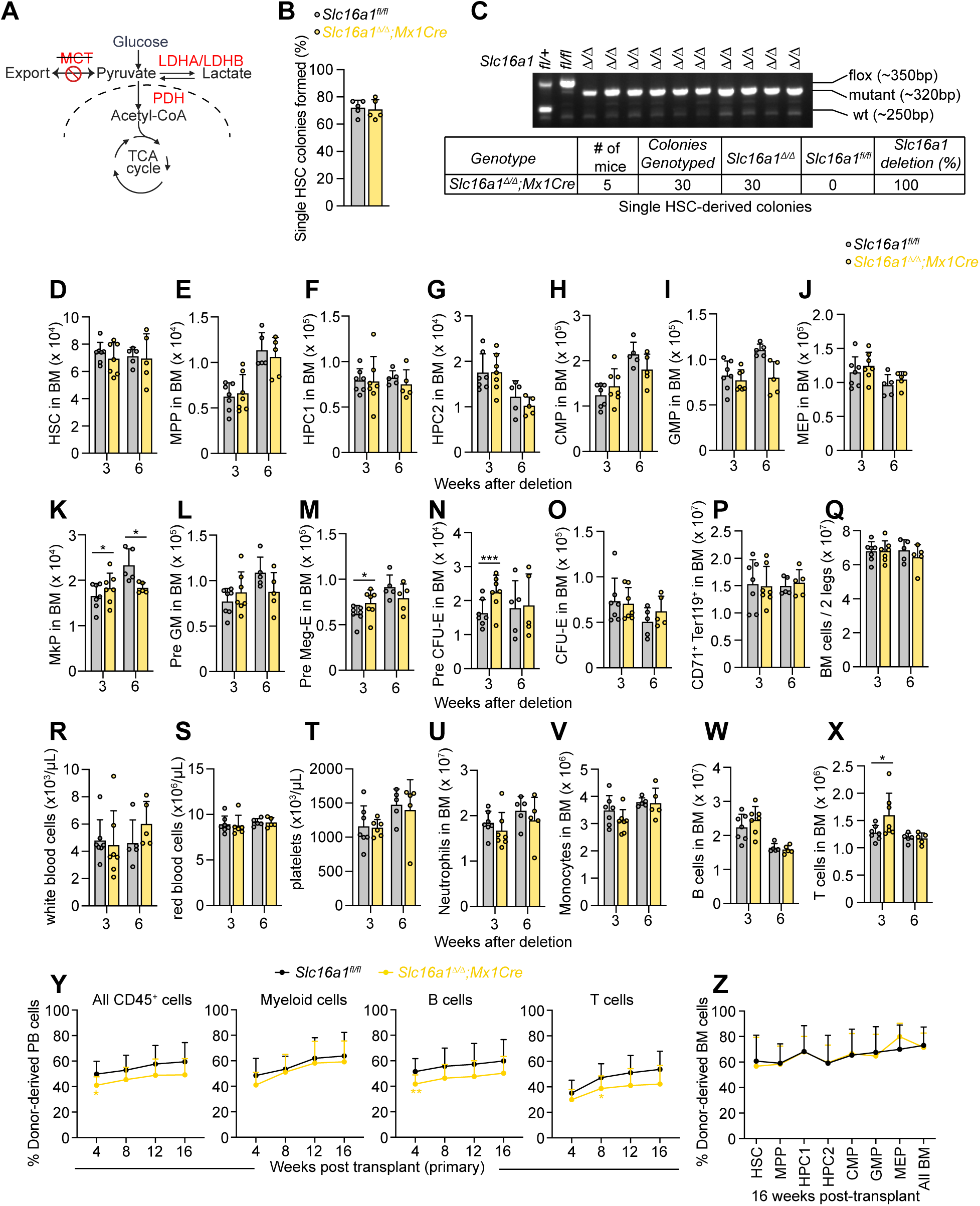
The effects of conditional *Slc16a1* deletion on hematopoiesis. **A,** Schematic of glycolysis termination illustrating conditional *Slc16a1* deletion with *Mx1Cre*. **B,** Colony-forming capacity in methylcellullose of single sorted *Slc16a1^Δ/Δ^* HSCs. **C,** Genotyping of colonies from **B** confirming 100% *Slc16a1* deletion in *Mx1Cre;Slc16a1^Δ/Δ^* HSCs. **D-P,** Numbers of HSCs, multipotent and restricted progenitors in the bone marrow of *Mx1Cre;Slc16a^Δ/Δ^* mice. **Q,** bone marrow cellularity in *Mx1Cre;Slc16a1^Δ/Δ^* mice. **R–T,** peripheral blood cell counts in *Mx1Cre;Slc16a1^Δ/Δ^* mice. **U–X,** numbers of neutrophils, monocytes, B, and T cells in *Mx1Cre;Slc16a1^Δ/Δ^*mice. **Y–Z,** Competitive bone marrow transplantation of donor CD45.2^+^ *Mx1Cre;Slc16a1^Δ/Δ^* or *Slc16a1^fl/fl^* littermate control bone marrow cells with wild-type CD45.1;CD45.2 competitor cells to lethally irradiated CD45.1 recipient mice. Shown are donor chimerism in **Y** peripheral blood at weeks 4-16 and **Z** BM HSPCs at week 16 (n = 13–14 mice per genotype for blood and BM; three independent experiments). All data represent mean ± s.d. Statistical significance was assessed with a t-test. All figures show * 0.01<p<0.05, ** 0.001<p<0.01, ***p<0.001.

**Figure S14.**
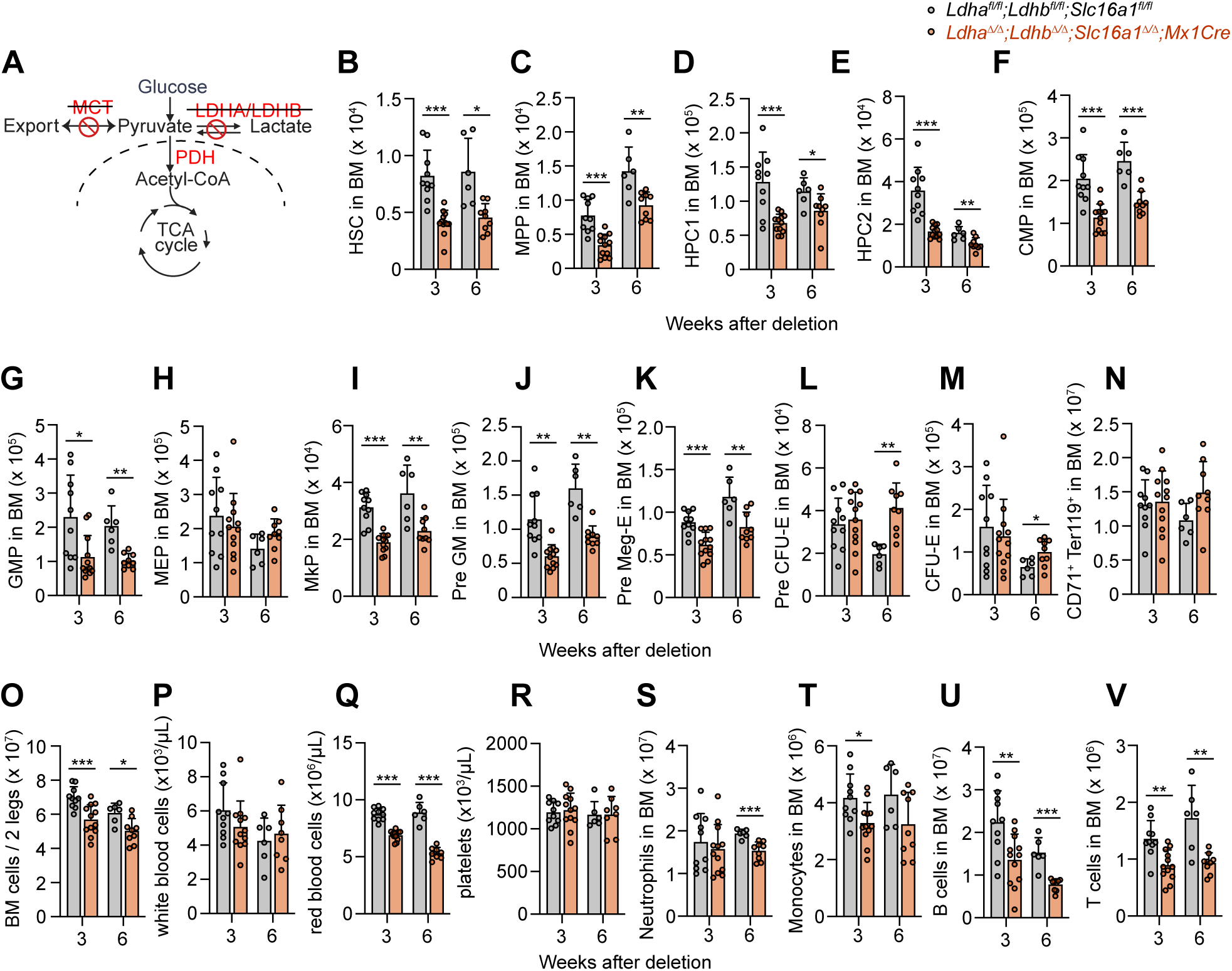
The effects of conditional *Ldha;Ldhb;Slc16a1* deletion on hematopoiesis. **A,** Schematic of glycolysis termination illustrating conditional *Ldha:Ldhb:Slc16a1* deletion with *Mx1Cre*. **B-N,** Numbers of HSCs, multipotent and restricted progenitors in the bone marrow of *Mx1Cre;Ldha^Δ/Δ^;Ldhb^Δ/Δ^*;*Slc16a1^Δ/Δ^* mice. **O,** bone marrow cellularity in *Mx1Cre;Ldha^Δ/Δ^;Ldhb^Δ/Δ^*;*Slc16a1^Δ/Δ^* mice. **P–R,** peripheral blood cell counts in *Mx1Cre;Ldha^Δ/Δ^;Ldhb^Δ/Δ^*;*Slc16a1^Δ/Δ^* mice. **S–V,** numbers of neutrophils, monocytes, B, and T cells in *Mx1Cre;Ldha^Δ/Δ^;Ldhb^Δ/Δ^*;*Slc16a1^Δ/Δ^* mice. All data represent mean ± s.d. Statistical significance was assessed with a Welch’s test (**B, G, J**) or a t-test (all others). All figures show * p<0.05, ** p<0.01, ***p<0.001.

**Figure S15.**
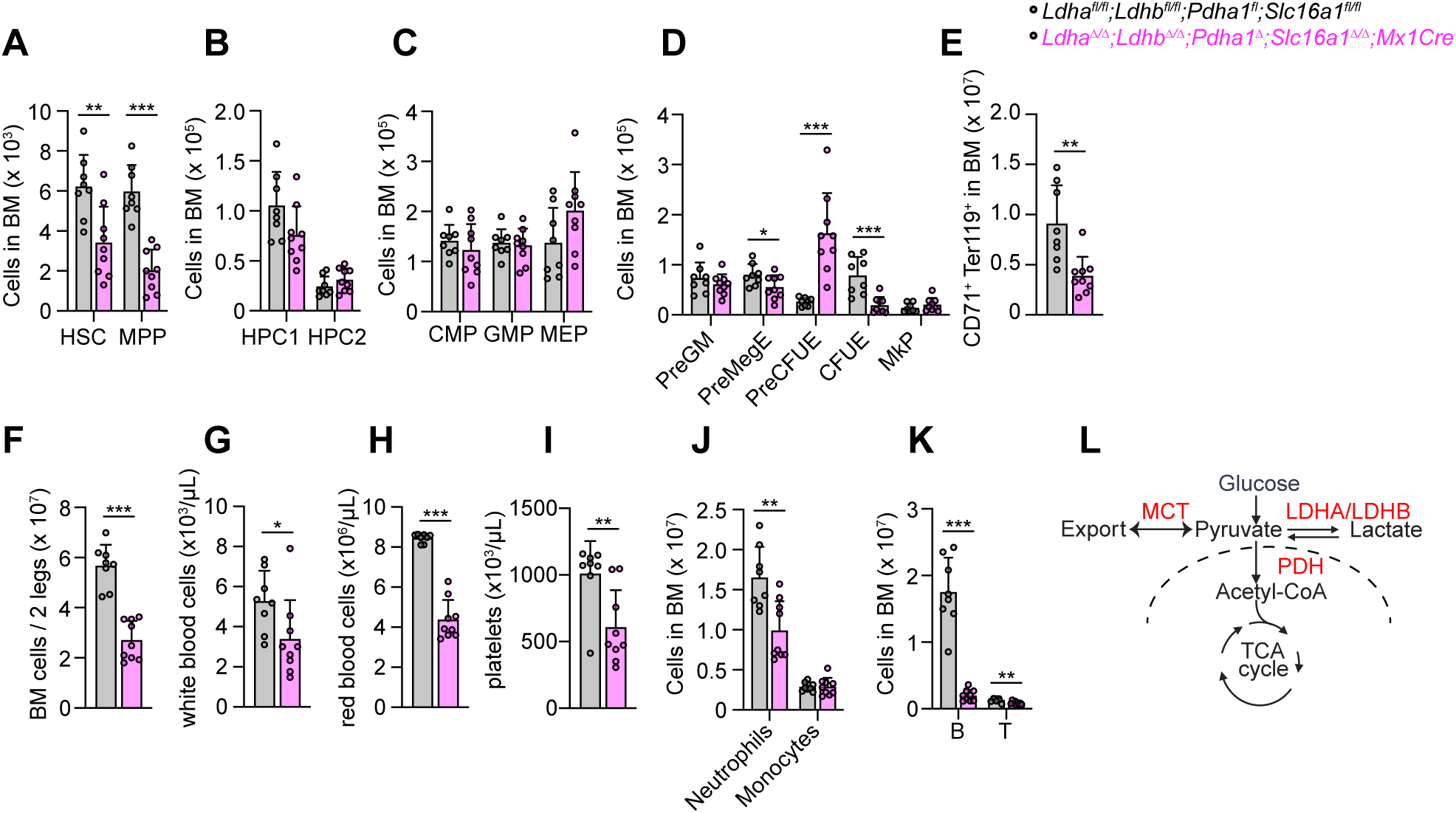
The effects of conditional *Ldha;Ldhb;Pdha1;Slc16a1* deletion on hematopoiesis. **A-E,** Numbers of HSCs, multipotent and restricted progenitors in the bone marrow of *Mx1Cre; Ldha^Δ/Δ^;Ldhb^Δ/Δ^*;*Pdha1^Δ^*;*Slc16a1^Δ/Δ^* mice 3 weeks after deletion. **F,** bone marrow cellularity in *Mx1Cre; Ldha^Δ/Δ^;Ldhb^Δ/Δ^*;*Pdha1^Δ^*;*Slc16a1^Δ/Δ^* mice. **G–I,** peripheral blood cell counts in *Mx1Cre; Ldha^Δ/Δ^;Ldhb^Δ/Δ^*;*Pdha1^Δ^*;*Slc16a1^Δ/Δ^* mice. **J–K,** numbers of neutrophils, monocytes, B, and T cells in *Mx1Cre; Ldha^Δ/Δ^;Ldhb^Δ/Δ^*; *Pdha1^Δ^*;*Slc16a1^Δ/Δ^*mice. **L,** Pyruvate export via MCTs is a third way to terminate glycolysis in addition to fermentation via LDH or oxidation via PDH. All data represent mean ± s.d. Statistical significance was assessed with a Mann-Whitney test (**G**, **I)**, Welch’s test (**J**) or a t-test (all others). All figures show * p<0.05, ** p<0.01, ***p<0.001

## References

1. Chandel, N.S. (2021). Glycolysis. Cold Spring Harb Perspect Biol 13. 10.1101/cshperspect.a040535.

2. Vander Heiden, M.G., Cantley, L.C., and Thompson, C.B. (2009). Understanding the Warburg effect: the metabolic requirements of cell proliferation. Science 324, 1029–1033. 10.1126/science.1160809.

3. Pasteur, L. (1861). Expériences et vues nouvelles sur la nature des fermentations. Comp. Rend. 52, 4.

4. Schwann, T. (1837). Vorläufige Mittheilung, betreffend Versuche über die Weingährung und Fäulniss. Annalen der Physik 11.

5. Meacham, C.E., DeVilbiss, A.W., and Morrison, S.J. (2022). Metabolic regulation of somatic stem cells in vivo. Nat Rev Mol Cell Biol 23, 428–443. 10.1038/s41580-022-00462-1.

6. Agathocleous, M., Love, N.K., Randlett, O., Harris, J.J., Liu, J., Murray, A.J., and Harris, W.A. (2012). Metabolic differentiation in the embryonic retina. Nat Cell Biol 14, 859–864. 10.1038/ncb2531.

7. Zheng, X., Boyer, L., Jin, M., Mertens, J., Kim, Y., Ma, L., Ma, L., Hamm, M., Gage, F.H., and Hunter, T. (2016). Metabolic reprogramming during neuronal differentiation from aerobic glycolysis to neuronal oxidative phosphorylation. Elife 5. 10.7554/eLife.13374.

8. Flores, A., Schell, J., Krall, A.S., Jelinek, D., Miranda, M., Grigorian, M., Braas, D., White, A.C., Zhou, J.L., Graham, N.A., et al. (2017). Lactate dehydrogenase activity drives hair follicle stem cell activation. Nat Cell Biol 19, 1017–1026. 10.1038/ncb3575.

9. Simsek, T., Kocabas, F., Zheng, J., Deberardinis, R.J., Mahmoud, A.I., Olson, E.N., Schneider, J.W., Zhang, C.C., and Sadek, H.A. (2010). The distinct metabolic profile of hematopoietic stem cells reflects their location in a hypoxic niche. Cell Stem Cell 7, 380–390. 10.1016/j.stem.2010.07.011.

10. Takubo, K., Nagamatsu, G., Kobayashi, C.I., Nakamura-Ishizu, A., Kobayashi, H., Ikeda, E., Goda, N., Rahimi, Y., Johnson, R.S., Soga, T., et al. (2013). Regulation of glycolysis by Pdk functions as a metabolic checkpoint for cell cycle quiescence in hematopoietic stem cells. Cell Stem Cell 12, 49–61. 10.1016/j.stem.2012.10.011.

11. Schell, J.C., Wisidagama, D.R., Bensard, C., Zhao, H., Wei, P., Tanner, J., Flores, A., Mohlman, J., Sorensen, L.K., Earl, C.S., et al. (2017). Control of intestinal stem cell function and proliferation by mitochondrial pyruvate metabolism. Nat Cell Biol 19, 1027–1036. 10.1038/ncb3593.

12. Luengo, A., Li, Z., Gui, D.Y., Sullivan, L.B., Zagorulya, M., Do, B.T., Ferreira, R., Naamati, A., Ali, A., Lewis, C.A., et al. (2021). Increased demand for NAD(+) relative to ATP drives aerobic glycolysis. Mol Cell 81, 691–707 e696. 10.1016/j.molcel.2020.12.012.

13. Wang, Y., Stancliffe, E., Fowle-Grider, R., Wang, R., Wang, C., Schwaiger-Haber, M., Shriver, L.P., and Patti, G.J. (2022). Saturation of the mitochondrial NADH shuttles drives aerobic glycolysis in proliferating cells. Mol Cell 82, 3270–3283 e3279. 10.1016/j.molcel.2022.07.007.

14. Ren, H., Tang, Y., and Zhang, D. (2025). The emerging role of protein L-lactylation in metabolic regulation and cell signalling. Nat Metab 7, 647–664. 10.1038/s42255-025-01259-0.

15. Sender, R., and Milo, R. (2021). The distribution of cellular turnover in the human body. Nat Med 27, 45–48. 10.1038/s41591-020-01182-9.

16. Jun, S., Mahesula, S., Mathews, T.P., Martin-Sandoval, M.S., Zhao, Z., Piskounova, E., and Agathocleous, M. (2021). The requirement for pyruvate dehydrogenase in leukemogenesis depends on cell lineage. Cell Metab 33, 1777–1792 e1778. 10.1016/j.cmet.2021.07.016.

17. Agathocleous, M., Meacham, C.E., Burgess, R.J., Piskounova, E., Zhao, Z., Crane, G.M., Cowin, B.L., Bruner, E., Murphy, M.M., Chen, W., et al. (2017). Ascorbate regulates haematopoietic stem cell function and leukaemogenesis. Nature 549, 476–481. 10.1038/nature23876.

18. Goldberg, E., Eddy, E.M., Duan, C., and Odet, F. (2010). LDHC: the ultimate testis-specific gene. J Androl 31, 86–94. 10.2164/jandrol.109.008367.

19. Jin, S., Chen, X., Yang, J., and Ding, J. (2023). Lactate dehydrogenase D is a general dehydrogenase for D-2-hydroxyacids and is associated with D-lactic acidosis. Nat Commun 14, 6638. 10.1038/s41467-023-42456-3.

20. Arafeh, R., Shibue, T., Dempster, J.M., Hahn, W.C., and Vazquez, F. (2025). The present and future of the Cancer Dependency Map. Nat Rev Cancer 25, 59–73. 10.1038/s41568-024-00763-x.

21. Zdralevic, M., Brand, A., Di Ianni, L., Dettmer, K., Reinders, J., Singer, K., Peter, K., Schnell, A., Bruss, C., Decking, S.M., et al. (2018). Double genetic disruption of lactate dehydrogenases A and B is required to ablate the “Warburg effect” restricting tumor growth to oxidative metabolism. J Biol Chem 293, 15947–15961. 10.1074/jbc.RA118.004180.

22. Guyon, J., Fernandez-Moncada, I., Larrieu, C.M., Bouchez, C.L., Pagano Zottola, A.C., Galvis, J., Chouleur, T., Burban, A., Joseph, K., Ravi, V.M., et al. (2022). Lactate dehydrogenases promote glioblastoma growth and invasion via a metabolic symbiosis. EMBO Mol Med 14, e15343. 10.15252/emmm.202115343.

23. Chen, X., Liu, L., Kang, S., Gnanaprakasam, J.R., and Wang, R. (2023). The lactate dehydrogenase (LDH) isoenzyme spectrum enables optimally controlling T cell glycolysis and differentiation. Sci Adv 9, eadd9554. 10.1126/sciadv.add9554.

24. Takubo, K., Goda, N., Yamada, W., Iriuchishima, H., Ikeda, E., Kubota, Y., Shima, H., Johnson, R.S., Hirao, A., Suematsu, M., and Suda, T. (2010). Regulation of the HIF-1alpha level is essential for hematopoietic stem cells. Cell Stem Cell 7, 391–402. 10.1016/j.stem.2010.06.020.

25. Nombela-Arrieta, C., Pivarnik, G., Winkel, B., Canty, K.J., Harley, B., Mahoney, J.E., Park, S.Y., Lu, J., Protopopov, A., and Silberstein, L.E. (2013). Quantitative imaging of haematopoietic stem and progenitor cell localization and hypoxic status in the bone marrow microenvironment. Nat Cell Biol 15, 533–543. 10.1038/ncb2730.

26. Spencer, J.A., Ferraro, F., Roussakis, E., Klein, A., Wu, J., Runnels, J.M., Zaher, W., Mortensen, L.J., Alt, C., Turcotte, R., et al. (2014). Direct measurement of local oxygen concentration in the bone marrow of live animals. Nature 508, 269–273. 10.1038/nature13034.

27. Wang, Y.H., Israelsen, W.J., Lee, D., Yu, V.W.C., Jeanson, N.T., Clish, C.B., Cantley, L.C., Vander Heiden, M.G., and Scadden, D.T. (2014). Cell-state-specific metabolic dependency in hematopoiesis and leukemogenesis. Cell 158, 1309–1323. 10.1016/j.cell.2014.07.048.

28. Guo, B., Huang, X., Lee, M.R., Lee, S.A., and Broxmeyer, H.E. (2018). Antagonism of PPAR-gamma signaling expands human hematopoietic stem and progenitor cells by enhancing glycolysis. Nat Med 24, 360–367. 10.1038/nm.4477.

29. Miharada, K., Karlsson, G., Rehn, M., Rorby, E., Siva, K., Cammenga, J., and Karlsson, S. (2011). Cripto regulates hematopoietic stem cells as a hypoxic-niche-related factor through cell surface receptor GRP78. Cell Stem Cell 9, 330–344. 10.1016/j.stem.2011.07.016.

30. 30. Cabezas-Wallscheid, N., Klimmeck, D., Hansson, J., Lipka, D.B., Reyes, A., Wang, Q., Weichenhan, D., Lier, A., von Paleske, L., Renders, S., et al. (2014). Identification of regulatory networks in HSCs and their immediate progeny via integrated proteome, transcriptome, and DNA methylome analysis. Cell Stem Cell 15, 507–522. 10.1016/j.stem.2014.07.005.

31. Vannini, N., Girotra, M., Naveiras, O., Nikitin, G., Campos, V., Giger, S., Roch, A., Auwerx, J., and Lutolf, M.P. (2016). Specification of haematopoietic stem cell fate via modulation of mitochondrial activity. Nat Commun 7, 13125. 10.1038/ncomms13125.

32. Qiu, J., Gjini, J., Arif, T., Moore, K., Lin, M., and Ghaffari, S. (2021). Using mitochondrial activity to select for potent human hematopoietic stem cells. Blood Adv 5, 1605–1616. 10.1182/bloodadvances.2020003658.

33. Mantel, C.R., O’Leary, H.A., Chitteti, B.R., Huang, X., Cooper, S., Hangoc, G., Brustovetsky, N., Srour, E.F., Lee, M.R., Messina-Graham, S., et al. (2015). Enhancing Hematopoietic Stem Cell Transplantation Efficacy by Mitigating Oxygen Shock. Cell 161, 1553–1565. 10.1016/j.cell.2015.04.054.

34. Watanuki, S., Kobayashi, H., Sugiura, Y., Yamamoto, M., Karigane, D., Shiroshita, K., Sorimachi, Y., Fujita, S., Morikawa, T., Koide, S., et al. (2024). Context-dependent modification of PFKFB3 in hematopoietic stem cells promotes anaerobic glycolysis and ensures stress hematopoiesis. Elife 12. 10.7554/eLife.87674.

35. Liang, R., Arif, T., Kalmykova, S., Kasianov, A., Lin, M., Menon, V., Qiu, J., Bernitz, J.M., Moore, K., Lin, F., et al. (2020). Restraining Lysosomal Activity Preserves Hematopoietic Stem Cell Quiescence and Potency. Cell Stem Cell 26, 359–376 e357. 10.1016/j.stem.2020.01.013.

36. Li, Z., Munim, M.B., Sharygin, D.A., Bevis, B.J., and Vander Heiden, M.G. (2024). Understanding the Warburg Effect in Cancer. Cold Spring Harb Perspect Med. 10.1101/cshperspect.a041532.

37. DeVilbiss, A.W., Zhao, Z., Martin-Sandoval, M.S., Ubellacker, J.M., Tasdogan, A., Agathocleous, M., Mathews, T.P., and Morrison, S.J. (2021). Metabolomic profiling of rare cell populations isolated by flow cytometry from tissues. Elife 10. 10.7554/eLife.61980.

38. Sidhu, S., Gangasani, A., Korotchkina, L.G., Suzuki, G., Fallavollita, J.A., Canty, J.M., Jr., and Patel, M.S. (2008). Tissue-specific pyruvate dehydrogenase complex deficiency causes cardiac hypertrophy and sudden death of weaned male mice. Am J Physiol Heart Circ Physiol 295, H946–H952. 10.1152/ajpheart.00363.2008.

39. Randle, P.J. (1986). Fuel selection in animals. Biochem Soc Trans 14, 799–806. 10.1042/bst0140799.

40. Wang, X., Menezes, C.J., Jia, Y., Xiao, Y., Venigalla, S.S.K., Cai, F., Hsieh, M.H., Gu, W., Du, L., Sudderth, J., et al. (2024). Metabolic inflexibility promotes mitochondrial health during liver regeneration. Science 384, eadj4301. 10.1126/science.adj4301.

41. Merchant, S., Paul, A., Reyes, A., Cassidy, D., Leach, A., Kim, D., Muh, S., Grabowski, G., Hoxhaj, G., Zhao, Z., and Morrison, S.J. (2025). Different effects of fatty acid oxidation on hematopoietic stem cells based on age and diet. Cell Stem Cell 32, 263–275 e265. 10.1016/j.stem.2024.11.014.

42. Wang, N., Jiang, X., Zhang, S., Zhu, A., Yuan, Y., Xu, H., Lei, J., and Yan, C. (2021). Structural basis of human monocarboxylate transporter 1 inhibition by anti-cancer drug candidates. Cell 184, 370–383 e313. 10.1016/j.cell.2020.11.043.

43. Hong, C.S., Graham, N.A., Gu, W., Espindola Camacho, C., Mah, V., Maresh, E.L., Alavi, M., Bagryanova, L., Krotee, P.A.L., Gardner, B.K., et al. (2016). MCT1 Modulates Cancer Cell Pyruvate Export and Growth of Tumors that Co-express MCT1 and MCT4. Cell Rep 14, 1590–1601. 10.1016/j.celrep.2016.01.057.

44. Felig, P., Pozefsky, T., Marliss, E., and Cahill, G.F., Jr. (1970). Alanine: key role in gluconeogenesis. Science 167, 1003–1004. 10.1126/science.167.3920.1003.

45. Jha, M.K., Lee, Y., Russell, K.A., Yang, F., Dastgheyb, R.M., Deme, P., Ament, X.H., Chen, W., Liu, Y., Guan, Y., et al. (2020). Monocarboxylate transporter 1 in Schwann cells contributes to maintenance of sensory nerve myelination during aging. Glia 68, 161–177. 10.1002/glia.23710.

46. Kuhn, R., Schwenk, F., Aguet, M., and Rajewsky, K. (1995). Inducible gene targeting in mice. Science 269, 1427–1429. 10.1126/science.7660125.

47. Rodriguez, C.I., Buchholz, F., Galloway, J., Sequerra, R., Kasper, J., Ayala, R., Stewart, A.F., and Dymecki, S.M. (2000). High-efficiency deleter mice show that FLPe is an alternative to Cre-loxP. Nat Genet 25, 139–140. 10.1038/75973.

48. Oguro, H., Ding, L., and Morrison, S.J. (2013). SLAM family markers resolve functionally distinct subpopulations of hematopoietic stem cells and multipotent progenitors. Cell Stem Cell 13, 102–116. 10.1016/j.stem.2013.05.014.

49. Kiel, M.J., Yilmaz, O.H., Iwashita, T., Yilmaz, O.H., Terhorst, C., and Morrison, S.J. (2005). SLAM family receptors distinguish hematopoietic stem and progenitor cells and reveal endothelial niches for stem cells. Cell 121, 1109–1121. 10.1016/j.cell.2005.05.026.

50. Akashi, K., Traver, D., Miyamoto, T., and Weissman, I.L. (2000). A clonogenic common myeloid progenitor that gives rise to all myeloid lineages. Nature 404, 193–197. 10.1038/35004599.

51. Pronk, C.J., Rossi, D.J., Mansson, R., Attema, J.L., Norddahl, G.L., Chan, C.K., Sigvardsson, M., Weissman, I.L., and Bryder, D. (2007). Elucidation of the phenotypic, functional, and molecular topography of a myeloerythroid progenitor cell hierarchy. Cell Stem Cell 1, 428–442. 10.1016/j.stem.2007.07.005.

52. Li, Y., Hook, J.S., Ding, Q., Xiao, X., Chung, S.S., Mettlen, M., Xu, L., Moreland, J.G., and Agathocleous, M. (2023). Neutrophil metabolomics in severe COVID-19 reveal GAPDH as a suppressor of neutrophil extracellular trap formation. Nat Commun 14, 2610. 10.1038/s41467-023-37567-w.

53. Su, X., Lu, W., and Rabinowitz, J.D. (2017). Metabolite Spectral Accuracy on Orbitraps. Analytical Chemistry 89, 5940–5948. 10.1021/acs.analchem.7b00396.

